# Comparative analysis of chemical cross-linking mass spectrometry data indicates that protein STY residues rarely react with N-hydroxysuccinimide ester cross-linkers

**DOI:** 10.1101/2023.01.17.524485

**Authors:** Yong Cao, Xin-Tong Liu, Peng-Zhi Mao, Ching Tarn, Meng-Qiu Dong

**Author notes:** To whom correspondence should be addressed: Y. C., M.-Q. D.

## Abstract

Chemical cross-linking of proteins coupled with mass spectrometry (CXMS) has enjoyed growing popularity in biomedical research. Most CXMS experiments utilize cross-linkers based on N-hydroxysuccinimide (NHS) ester, which react selectively with the amine groups found on the free N-termini of proteins and on the side chain of lysine (K) residues. It is also reported that under certain conditions they can react with the hydroxyl groups of serine (S), threonine (T), and tyrosine (Y). Some of the popular cross-link search engines including MeroX and xiSearch set STY, in addition to K, as cross-linkable sites by default. However, to what extent NHS ester cross-linkers react with STY under the typical CXMS experimental conditions remains unclear, nor has the reliability of STY-cross-link identifications. Here, by setting amino acids with chemically inert side chains such as glycine (G), valine (V), and leucine (L) as cross-linkable sites, which serves as a negative control, we show that software-identified STY-cross-links are only as reliable as GVL-cross-links. This is true across different NHS ester cross-linkers including DSS, DSSO, and DSBU, and across different search engines including MeroX, xiSearch, and pLink. Using a published dataset originated from synthetic peptides, we demonstrate that STY-cross-links indeed have a high false discovery rate. Further analysis revealed that depending on the data and the CXMS search engine used to analyze the data, up to 65% of the STY-cross-links identified are actually K-K cross-links of the same peptide pairs, up to 61% are actually K-mono-links, and the rest tend to contain short peptides at high risk of false identification.

## Introduction

Chemical cross-linking of proteins coupled with mass spectrometry analysis (abbreviated as CXMS, XL-MS or CLMS) is a convenient and effective tool to obtain three-dimensional structural information of proteins and protein complexes. In CXMS a small chemical cross-linker, which typically contains two amine-reactive groups, is used to form a covalent linkage between a pair of closely spaced amino acid residues, usually lysine residues. After trypsin digestion and liquid chromatography-tandem mass spectrometry (LC-MS/MS) analysis, cross-linked peptide pairs are identified from the CXMS data using a software tool such as pLink^1-2^, XlinkX^3-4^, MeroX^5^, or xiSearch^6-7^. A pair of cross-linked peptides, or a cross-link hereinafter, can be mapped to their parent protein(s) to locate the residue pair, that is, the two residues that are covalently linked by the cross-linker. By applying a distance restraint to all the residue pairs identified, one can use CXMS data to help build or improve atomic models of proteins or protein complexes^8-10^. More recently, CXMS applications have been expanded to detecting protein dynamics^11-12^, visualizing the process of protein unfolding^13^, and mapping protein-protein interactomes ^14-16^.

A multitude of cross-linkers have been developed for CXMS to target different amino acids residues such as lysine^17-22^, arginine^23^, glutamic/aspartic acid^24-26^, tyrosine^27^, or cysteine^28^. The most commonly used ones are homo-bifunctional cross-linkers based on N-hydroxysuccinimide (NHS) esters, including BS^3^, DSS, DSSO, and DSBU^29^. NHS esters react with primary amines, which include the ε-NH_2_ of lysine and the α-NH_2_ of a protein N-terminus. It is demonstrated that at near physiological pH of 6.7 and 7.8, an NHS ester cross-linker is highly specific towards the amine groups of peptides; even after 24 hours, there is no detectable reaction products with the peptide hydroxyl group on serine (S), threonine (T), or tyrosine (Y) unless there is a histidine residue in the same peptide^30^.

With respect to which amino acids are considered cross-linkable by NHS ester cross-linkers, the current cross-link search engines have opted differently^1-7, 31-38^. pLink and XlinkX, for example, have only lysine (K) and protein N-termini in the default setting. These two search engines thus output typically only K-K cross-links. Here, for simplicity of expression, a K- or K-K cross-link refers to a cross-link of which either or both link sites are K or a protein N-terminus. Other search engines like MeroX and xiSearch set KSTY and protein N-termini as cross-linkable sites by default. Adding STY as cross-linkable sites increases the number of identified residue pairs, with STY-cross-links making up as much as 30% of the total ^6, 39^. It is recognized that adding STY as cross-linkable sites increases the search space, which could increase false positive matches. To counter this negative effect, xiSearch gives K-STY and STY-STY cross-links less weight than K-K cross-links, and MeroX prohibits STY-STY cross-link identifications.

Although setting STY as cross-linkable sites is a somewhat common practice, the reliability of identified STY-cross-links has not been established firmly. We thus investigated this issue by comparing the STY-cross-links with the chemically impossible GVL-cross-links identified by the same software from the same CXMS data. The GVL-cross-links identified served as a negative control to estimate the extent of false identifications. We found that regardless of the search engine used, the number and the quality score distribution of STY- and GVL-cross-links were all comparable, this was true for different datasets generated by different laboratories using different NHS ester cross-linkers. This strongly suggests that STY-cross-link identifications are unreliable. We further verified that for STY- and GVL-cross-links the actual false discovery rate (FDR) far exceeded the intended FDR threshold, and uncovered two major sources of misidentification. Reviewing the evidence from this and other studies, we conclude that NHS ester cross-linkers generate few, if any, STY-cross-links under typical CXMS conditions. Adding STY as cross-linkable sites is not beneficial and cross-link search engines need to improve link-site localization function before they can be used to analyze CXMS data of cross-linkers that lack amino acid specificity such as photoactivated cross-linkers.

## Materials and Methods

### 2.1 software

pLink 2.3.9, MeroX 2.0.1.4, xiSearch 1.7.6.3 conjugated with xiFDR 2.1.5.2 were used to identify cross-linked peptides.

The open search of pFind 3 (3.1.6) was used to identify mono-link peptides of DSS and Leiker.

PTMiner (1.2.6) was used to re-localize DSS and Leiker modification sites.

### 2.2 datasets used in this study

The experimental conditions and access IDs of the nine datasets used in this study are summarized in following table.

### 2.3 Search parameters

#### For cross-linked peptides identification

Precursor mass tolerance 5 ppm, fragment ion mass tolerance 20 ppm, fixed modification C+57.021 Da, variable modification M+15.995 Da, peptide length minimum 5 amino acids and maximum 60 amino acids per chain, peptide mass minimum 500 and maximum 6,000 Da per chain, enzyme trypsin, three missed cleavage sites were allowed.

#### FDR cutoff

For xiSearch, the results were filtered by applying a 5% FDR cutoff at the residue pair level, and “boost” was checked. For pLink and MeroX, the results were filtered using a 5% FDR cutoff at the peptide pair level.

#### For mono-link peptides identification

Precursor mass tolerance 20 ppm, fragment ion mass tolerance 20 ppm, peptide length minimum 6 amino acids and maximum 100 amino acids per chain. The mono-link of DSS or Leiker on protein N-terminal, K, S, T, Y, G, V, and L was selected for the variable box. The results were filtered by applying a 1% FDR cutoff at both the peptide and protein group levels.

#### For mono-link sites localization

the pFind search parameter file “pFind.cfg” and identification file “pFind.spectra” were imported to PTMiner. transfer FDR (threshold 1%) was used for the localization of modification sites, and sites with probability less than 0.5 were filtered out.

### 2.4 link site setting

|  |  | KSTY group | KGVL group |
| --- | --- | --- | --- |
| **MeroX** | Alpha sites | K{ | K{ |
|  | Beta sites | KSTY{ | KGVL{ |
| **xiSearch** | Alpha sites | K(0) N-terminal (0)<br>S(0.2) T(0.2) Y(0.2) | K(0) N-terminal (0)<br>G(0.2) V(0.2) L(0.2) |
|  | Beta sites | K(0) N-terminal (0)<br>S(0.2) T(0.2) Y(0.2) | K(0) N-terminal (0)<br>G(0.2) V(0.2) L(0.2) |
| **pLink 2** | Alpha sites | KSTY[ | KGVL[ |
|  | Beta sites | KSTY[ | KGVL[ |

\* [ {stand for the N terminal of protein.

## Results

### STY-cross-links identified from CXMS data appear as reliable as chemically impossible GVL-cross-links across different NHS ester cross-linkers

To assess the reliability of STY-cross-links in CXMS identification results, we adopted a strategy that has been used in phosphoproteomics to estimate the false localization rate of phosphosites by treating alanine, which cannot be phosphorylated, as if it could.^43^ Likewise, we treated glycine (G), valine (V), and leucine (L), which cannot be cross-linked as if they could, to estimate the fraction of falsely identified STY-cross-links. To start with, we selected three datasets, each representing a different group of NHS ester cross-linkers. The BS^3^-BSA dataset was generated using a non-cleavable cross-linker BS^3^; the DSSO-BSA dataset was generated using a gas-phase cleavable cross-linker DSSO; and the DSBU-SurA/OmpA dataset was generated using DSBU, whose spacer arm can be cleaved to some extent in the gas phase. All three cross-likers are used frequently, and all three datasets originated from low-complexity samples containing only one (BSA) or two (SurA/OmpA) proteins.

xiSearch and MeroX, which routinely set K, S, T and Y as cross-linkable sites, were used to search for cross-links. As a negative control, we set K, G, V and L as cross-linkable sites and repeated the search. At 5% FDR cutoff, we obtained in most cases similar numbers of STY-cross-links and GVL-cross-links (Figure 1, compare STY and GVL within each panel). On the BS^3^-BSA dataset, xiSearch identified even more GVL-GVL cross-links than STY-STY cross-links (Figure 1A, 23 *vs*. 7), and MeroX identified more K-GVL cross-links than K-STY cross-links (Figure 1D, 102 *vs*. 64). Since the number of STY-cross-links did not surpass that by random match as estimated by GVL-cross-links for BS3, DSSO and DSBU, the reliability of the identified STY-cross-links becomes doubtful.

**Figure 1.**
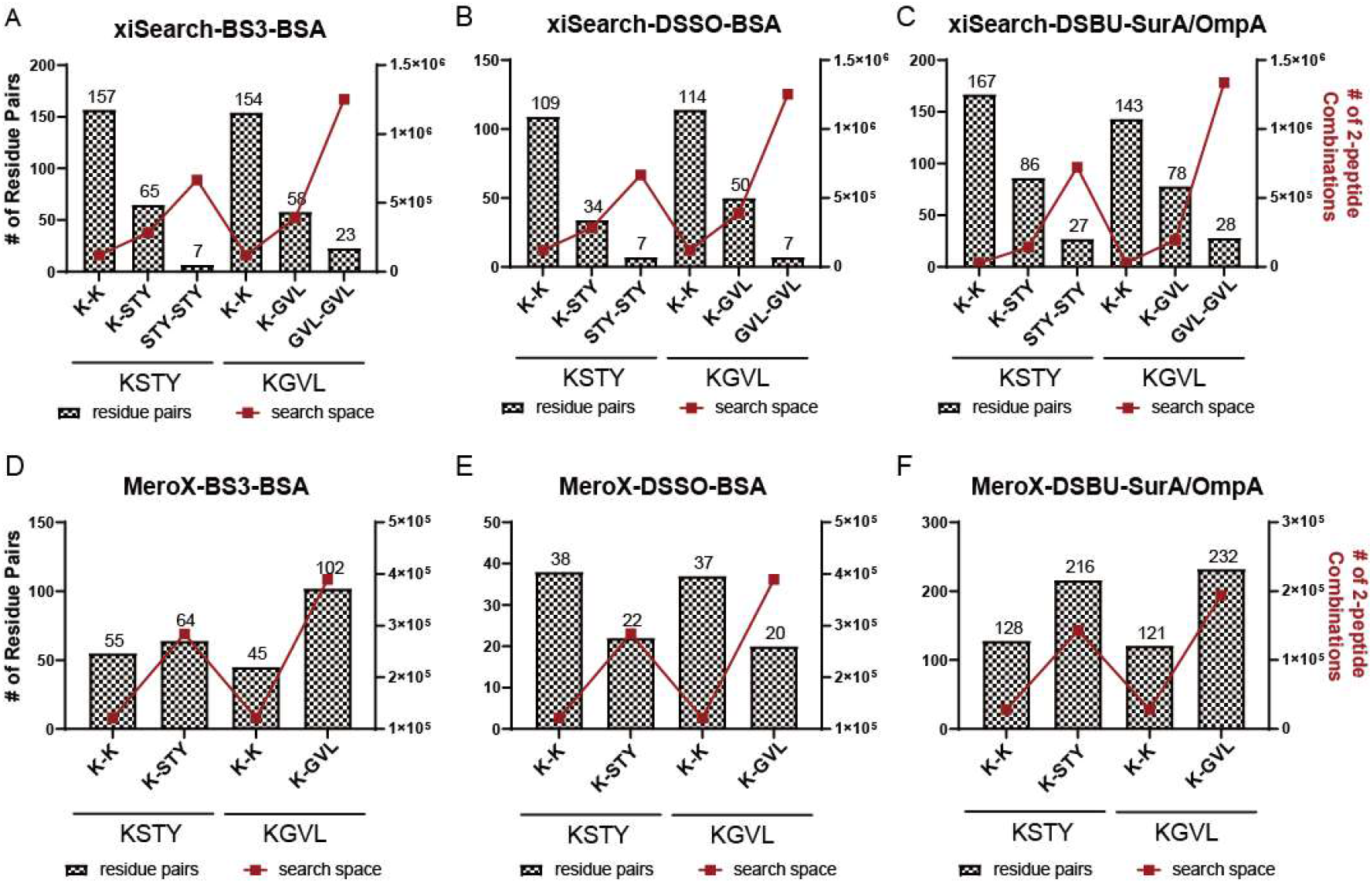
Similar numbers of STY-cross-links and GVL-cross-links were identified for different NHS ester cross-linker. (A-C) xiSearch identification. (D-F) MeroX identification. KSTY or KGVL were set as cross-linkable sites. The number of identified residue pair are indicated by bars (left y-axis) and the size of the search space as measured by the numbers of possible, link-site-sensitive peptide pair combinations is indicated by line-connected red squares (right y-axis).

### STY-cross-links identified using different cross-link search engines are similarly questionable

Wondering whether misidentification of STY-cross-links is a general problem in CXMS data analysis, we included another cross-link search engine pLink 2^2^ in further investigations. Two test datasets were used at this stage: the DSSO-Ribo dataset represents a median-complexity sample of purified *E. coli* ribosomes cross-linked with DSSO, and the DSS-Mpneu dataset from *Mycoplasma pneumoniae* cells cross-linked in-situ using DSS. On the DSSO-Ribo dataset, xiSearch, MeroX and pLink all produced more GVL-cross-links than STY-cross-links (Figure 2A-C). Take the xiSearch results as an example, setting KSTY or KGVL as cross-linkable sites yielded, respectively, 829 or 771 K-K cross-links, 95 or 151 K-nonK cross-links, and 7 or 30 nonK-nonK cross-links (Figure 2A). Similar observations were made on the DSS-Mpneu dataset using xiSearch and pLink 2 (Supplementary Figure 1A-B). We were not able to complete a MeroX search on large datasets like this one, so only xiSearch and pLink 2 search results were available for the high-complexity samples. In sum, regardless of sample complexity, cross-linker, and data analysis software, the number of STY-cross-links identified from any of the datasets were not above the number of GVL-cross-links identified from the same data. Therefore, judged by the number of identifications, STY-cross-links are as reliable as the chemically impossible GVL-cross-links.

**Figure 2.**
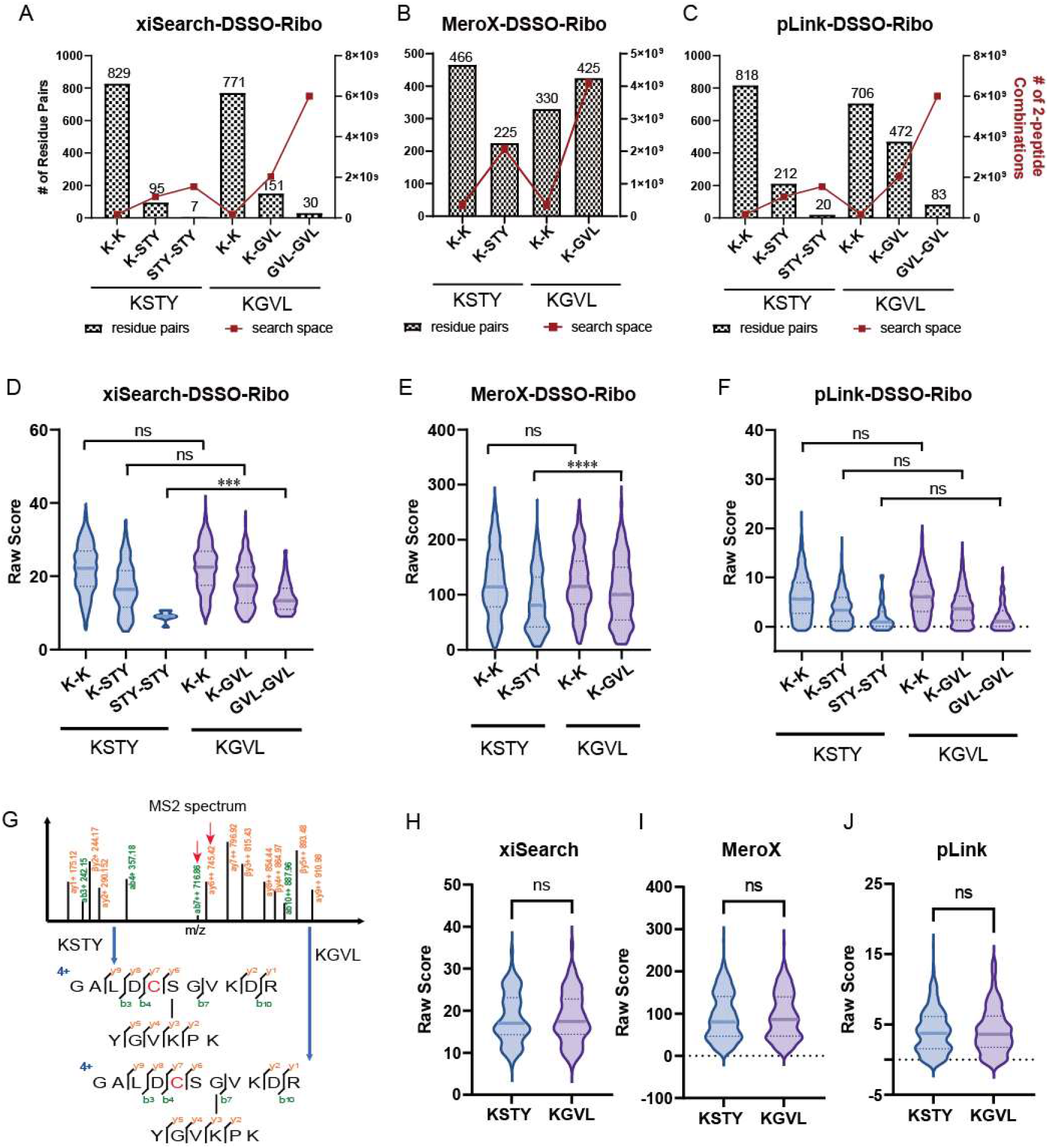
STY-cross-links are only as reliable as GVL-cross-links regardless of the search engines used to identify them. The numbers (A-C) and the CSM score distributions (D-F) of the STY- and GVL-cross-link identifications by xiSearch, MeroX, or pLink on the medium-complexity DSSO-Ribo dataset. In (A-C) the number of identified residue pair are indicated by bars (left y-axis) and the size of the search space as measured by the numbers of possible, link-site-sensitive peptide pair combinations is indicated by line-connected red squares (right y-axis). (G) Example of a dual-identify MS2 spectrum. (H-J) CSM score distribution of the dual-identity MS2 spectra found from the xiSearch (n=78, *p*=0.783), MeroX (n=252, *p*=0.258), and pLink (n=426, *p*=0.051).

Next, we asked whether the identified STY-cross-links scored better than the GVL-cross-links in cross-link-spectrum match (CSM). As shown in Figure 2D-F (DSSO-Ribo) and Supplementary Figure 1C-D (DSS-Mpneu), for all three search engines tested, the median CSM scores of the identified STY-cross-links are similar to, or lower than, those of the GVL-cross-links, the median CSM score of the K-K cross-links, as a reference, are always higher than those of the K-nonK and nonK-nonK cross-links identified from the same data; and there is no difference in the CSM score distribution of the K-K cross-links identified between the KSTY search and KGVL search (Figure 2D-F and Supplementary Figure 1C-D).

Additionally, we noticed that some of the MS2 spectra received dual identities, that is, they were identified as STY-cross-links in the KSTY search, and as GVL-cross-links in the KGVL search. One such example is shown in Figure 2G. As can be seen, a strong y_6_^++^ fragment ion and a weak b_7_^++^ fragment ion of the α-peptide GALDCSGVKDR (labeled as αy6++ 745.42 and αb7++ 716.65 on the spectrum) are able to position the DSS link site to -SG- in the middle of the α-peptide, but since there is no detected cleavage between S and G to pinpoint the link site, this spectrum is matched either as a S-cross-link in KSTY search or a G-cross-link in KGVL search. Intrigued by this, we collected all the dual-identity spectra found in this study and analyzed their CSM scores. As shown in Figure 2H-I, their CSM score distributions seem identical either as STY-cross-links or as GVL-cross-links, regardless of the search engine used. Therefore, judged by the number of identified cross-links as well as the quality of the CSMs, the STY-cross-links are not better than the GVL-cross-links, and all the search engines seem to share this problem.

### STY-mono-links identified using a PTM analysis workflow are also questionable

The formation of a cross-link involves two ligation reactions, one on each end of a cross-linker. More likely than not, the two ligation reactions occur one after another. The intermediate product—a mono-link—can go on to form a cross-link, or not if the remaining NHS ester is hydrolyzed. In the latter case, a mono-link becomes an end product. In our experience, there are more mono-links than cross-links^21^. We reasoned that if there are truly STY-cross-links, there should be STY-mono-links and they are likely more abundant.

Although many cross-link search engines output mono-link identifications, some such as MeroX do not. For this reason and that a mono-link is a linear peptide with a small chemical modification, we used Open-pFind^44^ to identify mono-links. Of the three datasets used, two originated from DSS-linked samples and one from a Leiker-linked sample. Leiker is a NHS ester cross-linker with a biotin tag^21^. In Open-pFind search, K, S, T, Y, G, V, and L were set as variable modification sites for DSS and Leiker, with 1% FDR cutoff at both the peptide level and the protein group level. Following re-evaluation and correction of the modification sites by PTMiner^45^. We obtained >7,000 DSS mono-links on K, and 4-17 times less DSS mono-links on STY or GVL (Figure 3). More importantly, the number of identified mono-links on STY were less than those on GVL, which cannot be modified. These results suggest that more likely than not, the STY-mono-links identified from samples treated with NHS ester cross-linker are spurious matches.

**Figure 3.**
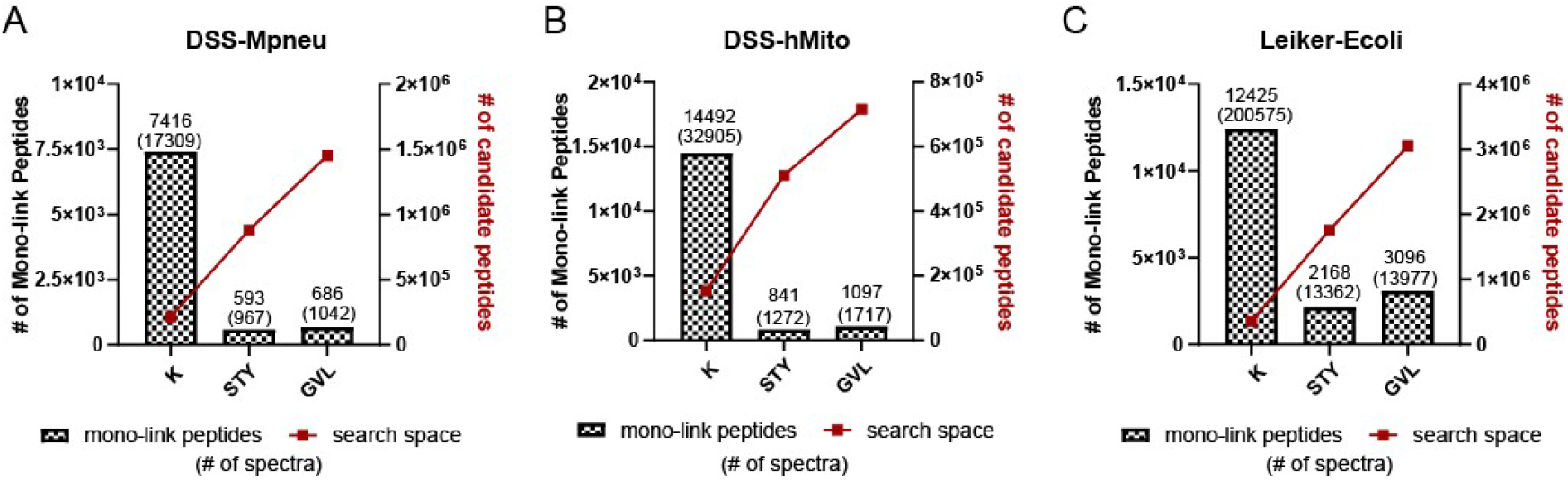
Mono-links of NHS ester cross-linkers identified by Open-pFind. The bars indicate the numbers of mono-link peptides for DSS (A-B) or Leiker (C) from the indicated datasets (left y-axis), with the number of corresponding MS2 spectra shown in parentheses, and the number of candidate peptides indicated by line-connected red squares (right y-axis).

In agreement with the above conclusion, a recent proteome-scale study has found no evidence of a NHS ester probe targeting S, T, or Y^46^. In this study, a HeLa cell lysate was incubated with a pair of light and heavy isotope encoded NHS ester probe, and pChem—a software tool developed specifically to assign modification sites of chemical probes—was employed to analyze the data. The authors found that the NHS ester probe modified only lysine residues and protein N-termini.

Taken together, we conclude that under the conditions typically used in CXMS experiments, it should be rare for an NHS ester cross-link to react with S, T, or Y, if ever it does. The STY-cross-links identified by the current versions of cross-link search engines are mostly unreliable.

### Setting STY as additional cross-linkable sites greatly increases FDR

To quantify the effect of setting STY as additional cross-linkable sites on cross-link identifications, we turned to the benchmark datasets of synthetic peptides^42^. To generate these datasets, 95 chemically synthesized peptides of a Cas9 protein were divided into 12 group, cross-linked within-group and pooled together for LC-MS/MS after the cross-linking reactions were stopped. Thus, between-group cross-links, if identified, must be false; whereas within-group cross-links may be true. This provides a means to estimate FDR experimentally, independent of the software estimated FDR. because some of the within-group cross-link identifications may be false, this experimental FDR, calculated by dividing the number of between-group cross-links with the sum of between- and within-group cross-links, estimates the lower boundary of the actual FDR, We used two benchmark dataset, generated with DSS and another with DSBU. The MS data were searched with or without setting STY as cross-linkable sites in addition to K and protein N-termini. A cutoff of 5% software estimated FDR was applied at the peptide pair level. The K-K, K-STY, and STY-STY cross-link identifications from the KSTY search were examined separately. As shown in Figure 4, for all three search engines tested and xiSearch in particular, the STY-cross-links identified had much higher experimental FDR than the K-K cross-links identified either from a KSTY search or a K only search. The experimental FDR of the STY-STY-cross-links identified by xiSearch reached up to 100% (Figure 4A and 4D). Invariably, KSTY search increased experimental FDR and slightly decreased the number of legitimate K-K cross-link identifications. This result shows that setting STY as additional cross-linkable sites brings more harm than benefit.

**Figure 4.**
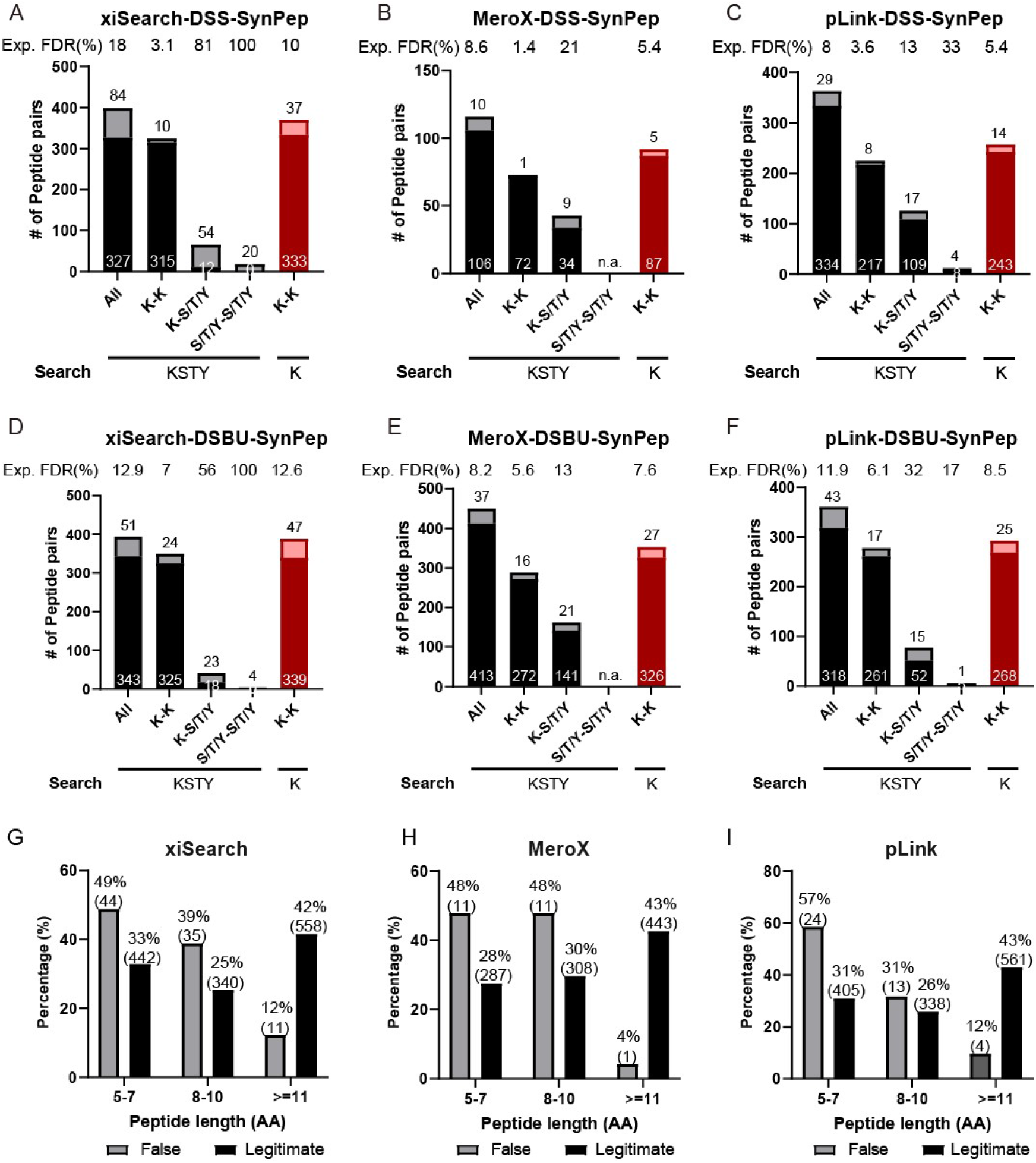
Experimental FDR of STY-cross-links estimated using two benchmark datasets of synthetic peptides. (A-C) Search results of the DSS-SynPep dataset by xiSearch, MeroX, and pLink. (D-F) Search results of the DSBU-SynPep dataset by xiSearch, MeroX, and pLink. (G-I) Distribution of the length of the peptides constituting false cross-links versus those of the legitimate cross-links identified by the indicated search engine. Note: these two datasets were searched against cas9 and 293 common contaminant protein sequences. The identification results were filtered using a 5% software estimated FDR at the peptide pair level. Light shade denotes the false between-group cross-links and dark shade denotes the legitimate, within-group cross-links. In (G-I), the number of peptides in each bin is shown in parentheses.

We also analyzed the distribution of the length of the peptides constituting the false, between-group cross-links. Compared with the legitimate cross-links, the false ones tend to have short peptides of 5-7 aa (Figure 4 G-H). The implication of this result will be discussed later.

### A subset of STY- or GVL-cross-links are misidentified K-K cross-links

Having found that most if not all STY-cross-link identifications are probably false, we asked what might have led to the erroneous identifications. A close examination of the problematic MS2 spectra revealed that a subset of them are actually K-K cross-links misidentified as STY- or GVL-cross-links.

One example is shown in Figure 5A-E. A 5^+^ precursor ion whose monoisotopic peak is *m/z* 576.596 is a cross-link between DRVEDATLVLSVGDEVEAKFTGVDRK (α peptide) and KGALVTGK (β peptide). Across the chromatogram peak, which apexed at 74.2 min and trailed to 76 min, four MS2 spectra were acquired and identified respectively as K^α19^-K^β1^, V^α23^-K^β1^, K^α19^-G^β2^, and K^α19^-K^β1^ cross-links. The first (Figure 5B) and the fourth (Figure 5E) MS2 spectra both have contiguous b- and y-series to pinpoint the link sites to K^α19^ and K^β1^. The second MS2 has two tiny peaks at *m/z* 628.35 and *m/z* 908.43 that happen to match the theoretical *m/z* of αSy_5_^+^ and αLy ^+^, respectively (Figure 5C). As a result, the link site on the α peptide is assigned to V^α23^, instead of K^α19^. The third MS2 (Figure 5D) is of the lowest base peak intensity (2.4e+04), compared to the other three (2.7e+04 to 5.2e+05). The *m/z* 129.10 ion, interpreted as βb_1_^+^, is the reason to localize the link site on the β-peptide to G^β2^ instead of K^β1^. However, according to the peptide fragmentation mechanism, which is deduced from numerous experimental observations and validated by theoretical calculations, there is no b1 ion under normal circumstances. Another and better interpretation of *m/z* 129.10 is [y_1_-H_2_O]^+^ from either α- or β-peptide, as both of them have a C-terminal lysine residue. When the same MS2 data were searched again by specifying K instead of KGVL as cross-link sites, all four of them were identified as K^α19^-K^β1^ cross-links of the same peptide pair.

**Figure 5.**
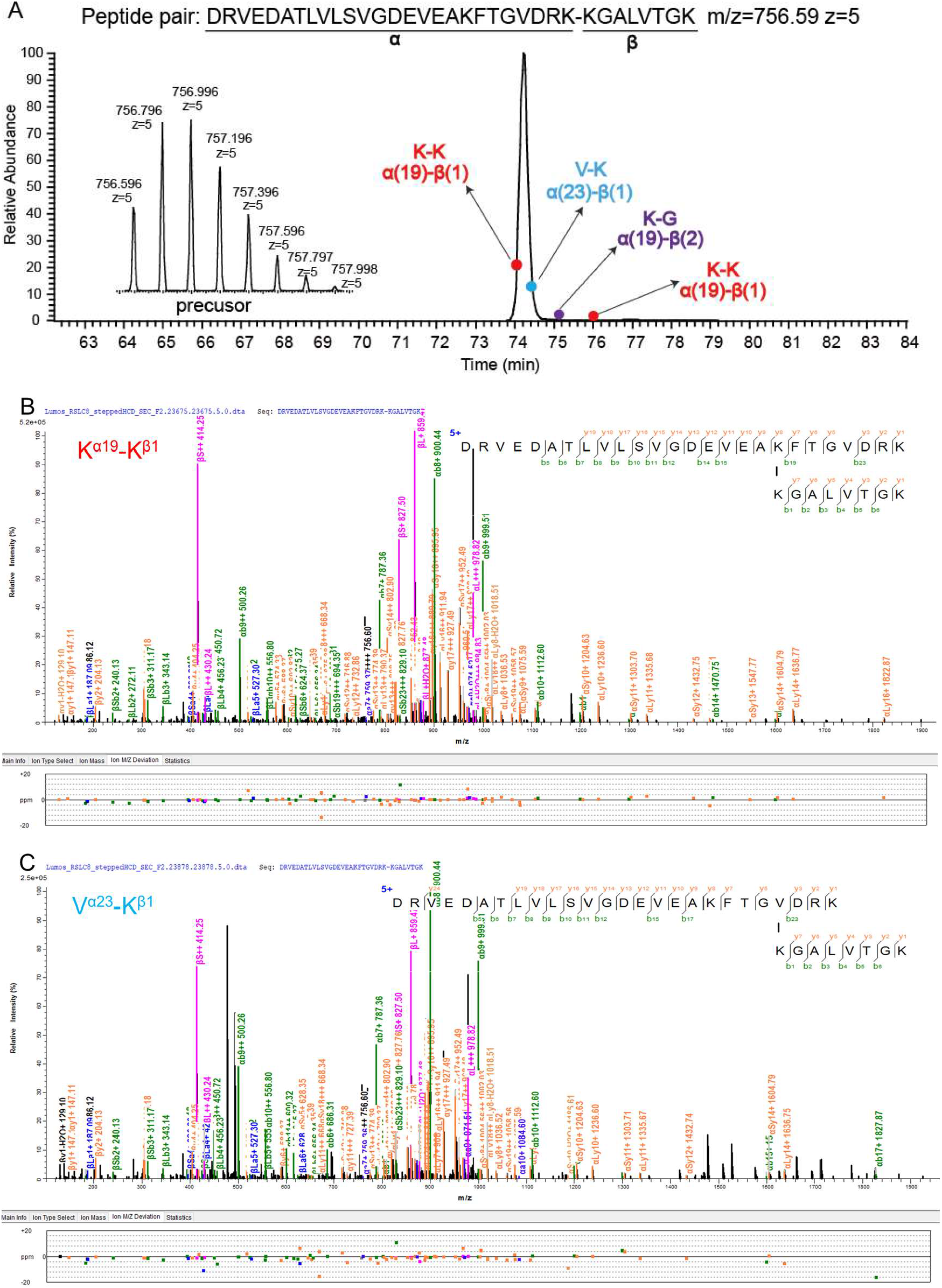
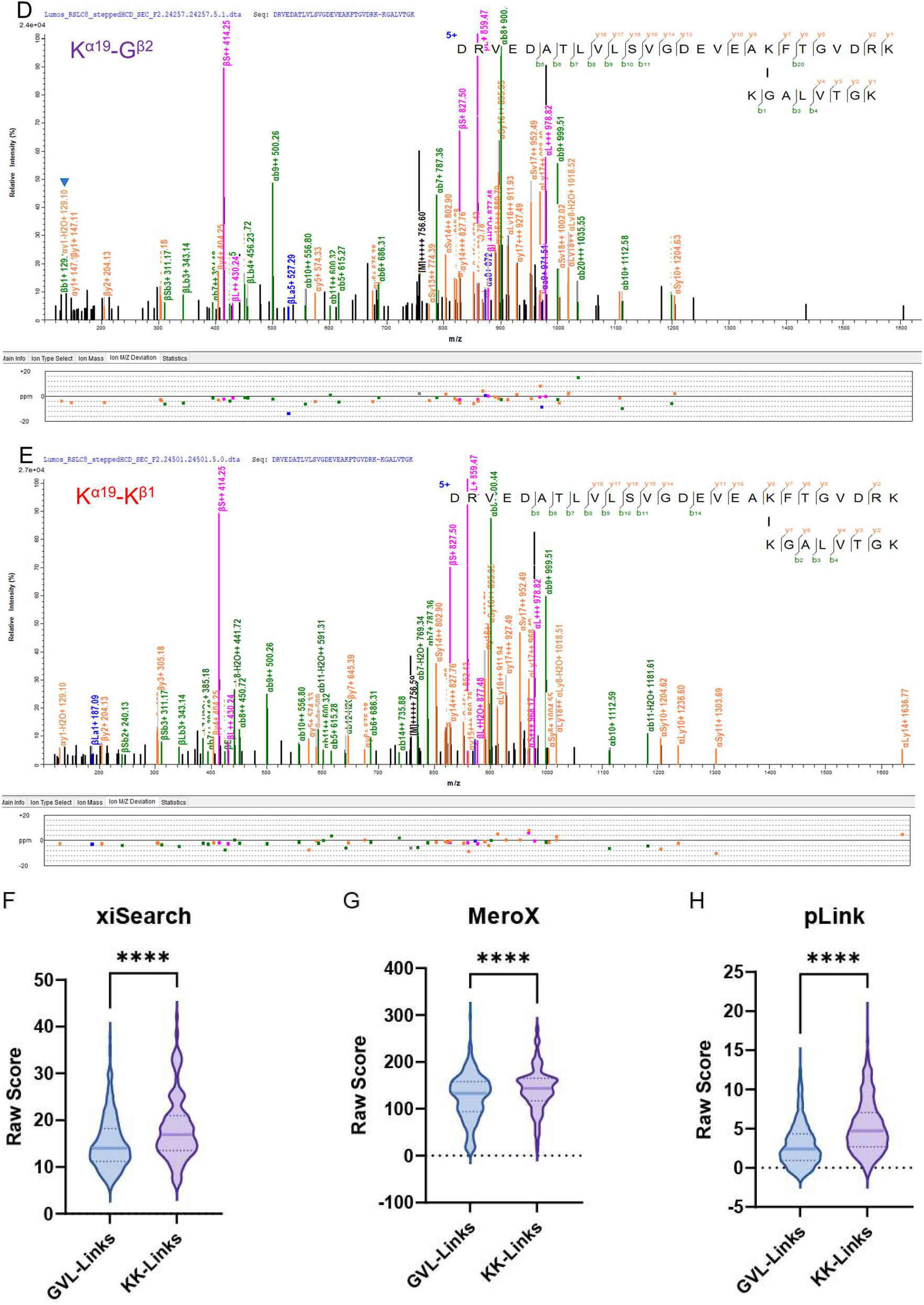
K-K cross-links misidentified as GVL-cross-links. (A) The extracted ion chromatogram (XIC) of a pair of cross-linked peptides DRVEDATLVLSVGDEVEAKFTGVDRK-KGALVTGK (*m/z*=756.59 z=5). Three link-site isoforms were identified by four MS2 spectra across the chromatogram peak. Shown on the left is the isotopic peak cluster of the precursor ion. (B-E) The four MS2 spectra of the cross-linked peptide pair in (A), annotated according to their link-site identifications. (F-H) Comparison of the scores between the GVL-cross-links and their cognate K-K cross-links identified from the same data by xiSearch (n =283, *p*<0.0001), MeroX (n =218, *p*<0.0001), and pLink (n =768, *p*<0.0001).

In total, we found from the xiSearch, MeroX, and pLink search results 283, 218, 768 pairs of apparent link site isoforms, respectively. In each pair are two cross-links of the same two peptides, identified from the same raw file, but one is a K-K cross-link and the other is a GVL-cross-link. Paired comparison of the best MS2 of the K-K cross-link and the best MS2 of the GVL-cross-link finds that the former has a higher CSM score than the latter in most cases (*p*<0.0001) (Figure 5F-H). Naturally, the MS2 spectra of such GVL-cross-links, if searched again with K instead of KGVL as the cross-linkable link site, are then identified as their cognate K-K cross-links. (Supplementary Table 5).

Similar observations were made on the STY-cross-links. In the example shown in Figure 6A-C, a cross-link between PWNSTWFANTKEFADNLDSDFK (α-peptide) and LVLERPAKSL (β-peptide) was identified twice across the chromatogram peak of the 4^+^ precursor at monoisotopic *m/z* 1018.752 (Figure 6A). However, the link sites were identified as K^α11^-K^β8^ at the apex from a high signal intensity MS2 (base peak intensity 1.3e+06, Figure 6C) and as T^α10^-S^β9^ in the ascending phase of from a lower intensity MS2 (1.7e+05, Figure 6B). The higher-but not the lower-intensity MS2 has fragment ions to pinpoint the link site on the α-peptide. On the β-peptide, assigning the link site to S^β9^ instead of K^β8^ relies entirely on a tiny peak of *m/z* 429.24, which happens to match the theoretical *m/z* of βSy_3_ ^+^ (Figure 6B). When the cross-linkable site was set to K, this MS2 spectrum (Figure 6B) was identified as K^α11^-K^β8^ cross-link, just like the other one (Figure 6C).

**Figure 6.**
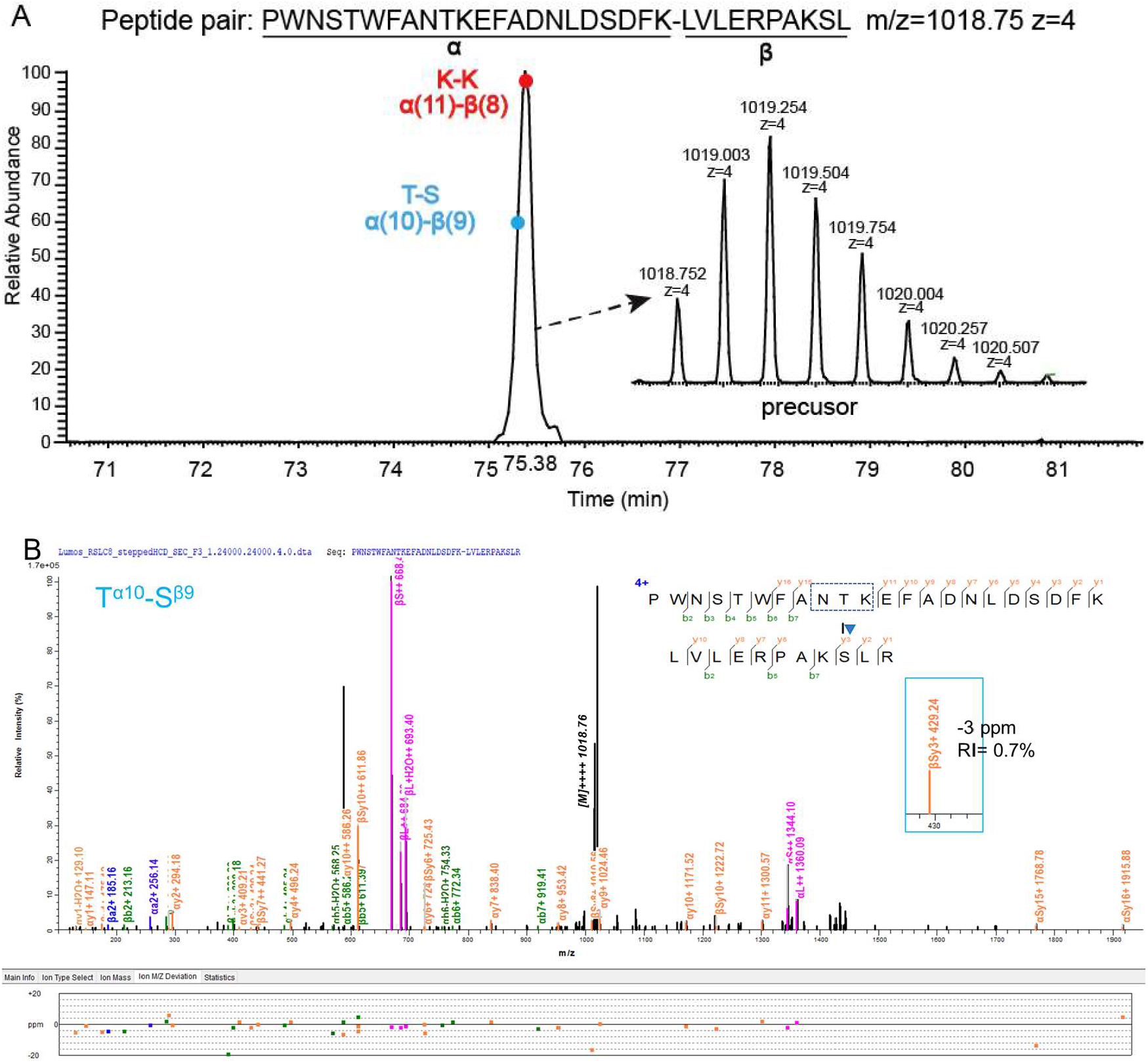
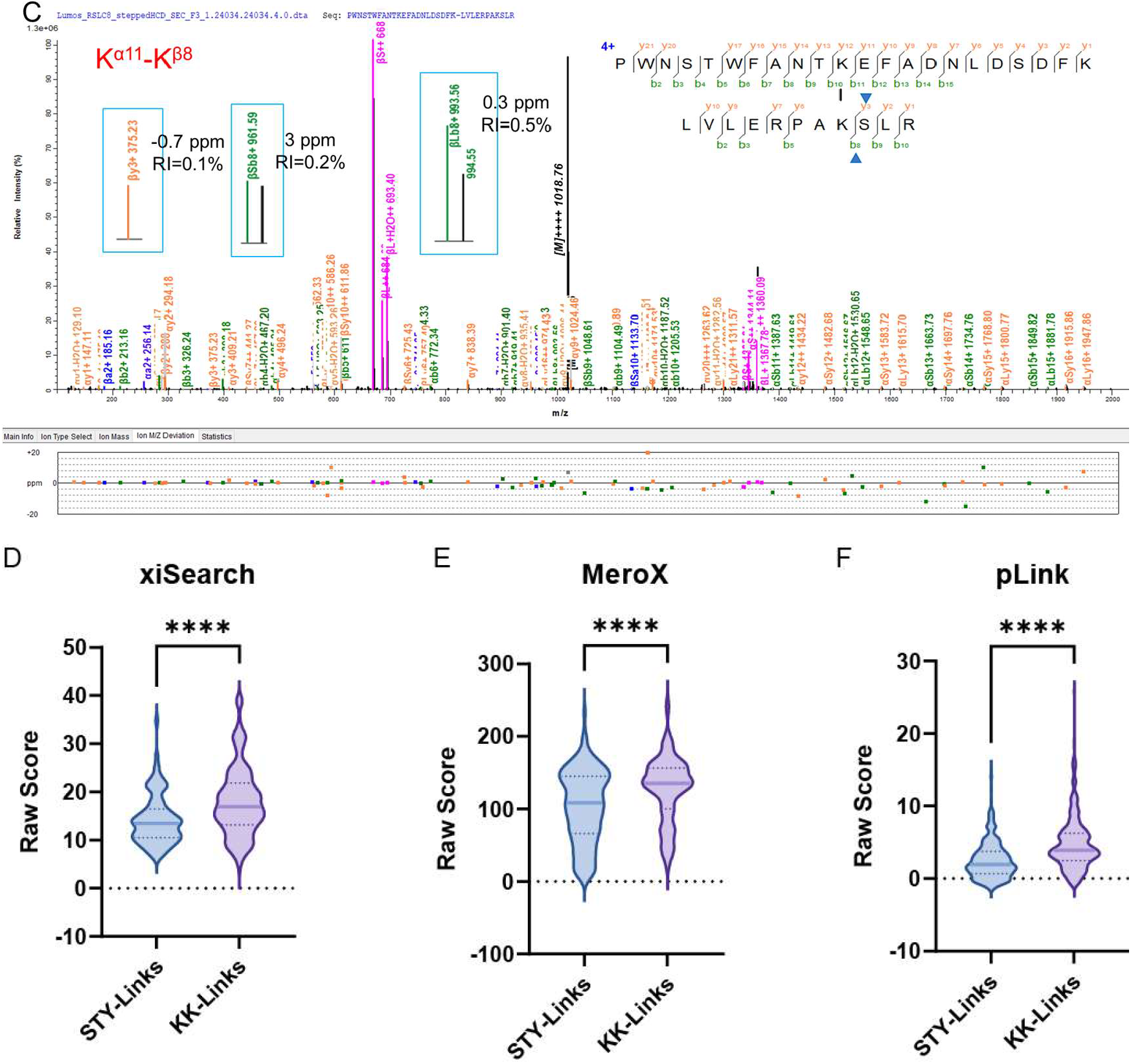
K-K cross-links misidentified as STY-cross-links. (A) The XIC of a pair of cross-linked peptides SGKSELEAFEVALENVRPTVEVK-VKHPSELVNVGDELTVK (*m/z*=1018.75 z=4). Two link-site isoforms were identified from the indicated positions in XIC (solid circles). The isotopic peak cluster of the precursor was shown on the right. (B-C) The MS2 spectra of two link-site isoform, T^α10^-S^β9^ and K^α11^-K^β8^. (RI=relative intensity) (D-F) Comparison of the scores between the GVL-cross-links and their cognate K-K cross-links identified from the same data by xiSearch (n =175, *p*<0.0001), MeroX (n =150, *p*<0.0001), and pLink (n =459, *p*<0.0001).

Supplementary Figure 3 shows another example. Cross-links between SGKSELEAFEVALENVRPTVEVK (α-peptide) and VKHPSELVNVGDELTVK (β-peptide) were identified. MS2 from the 4^+^ precursor was identified as a STY-cross-link (S^α1^-K^β2^), while that from 5^+^ precursor was identified as a K-K cross-link (K^α3^-K^β2^) (Supplementary Figure 3). The MS2 of the 5^+^ precursor, in contrast, is a high-quality spectrum with contiguous fragment ion series to pinpoint the link site to K^α3^ (Supplementary Figure 3B). In contrast, the MS2 of the 4^+^ precursor is a poor-quality spectrum with many missing ions (Supplementary Figure 3C). No fragment ions are present to pinpoint the link site on the α-peptide; it could be any one of the first nine residues if the NHS ester cross-linking chemistry is ignored. Therefore, we conclude that the S^α1^-K^β2^ cross-link is identified incorrectly, it is actually a K-K cross-link.

In the xiSearch, MeroX, and pLink search results, we found a total of 175, 150, and 459 MS2 spectra that behave like this, that is, when the cross-linkable sites are set differently, the peptide sequence identifications remain the same, but link site identifications do not. Paired comparison finds that the K-K cross-links have higher CSM scores than the STY-cross-links in most cases (*p*<0.0001, Figure 6D). This indicates that some of the STY-cross-links are in fact K-K cross-links. Similar with the GVL-cross-links, the MS2 spectra of such STY-cross-links, if searched again with K-, not KSTY-, as cross-linkable sites, are identified as their cognate K-K cross-links. (Supplementary Table 5).

Below is a brief account of our findings from inspecting the MS2 spectra of questionable cross-link identifications. In some of the MS2 spectra we found no ions supporting a STY/GVL link site due to the absence of certain cleavage products (Figure 6B). For others, it is often clear that their assumed identities are dubious as they have the following characteristics.

1. Unaccompanied by isotopic peaks to verify their assumed charge state (Supplementary Figure 4A-B).
2. Low intensity—often not above the tiny peaks at the “grass” level (Supplementary Figure 4C).
3. Large mass deviation—often an outlier relative to the mass deviations of other fragments ions (Supplementary Figure 4B S4D).
4. Can be interpreted differently, e.g., as an isotopic peak of a different fragment ion (Supplementary Figure 4E). *m/z* 129.10 is a frequent “offender” in this category. It is [y_1_-H_2_O]^+^ when a peptide has a K at the C-terminus, but when interpreted by the search engine as b_1_^+^ for a peptide with K at the N-terminus, it becomes evidence against assigning the link site to the N-terminal K. However, the latter interpretation is incorrect because peptides normally do not produce b_1_^+^ ions^47^. Examples are shown in Supplementary Figure 4E in addition to Figure 5, Figure 6, and Supplementary Figure 3.

### A subset of STY- or GVL-cross-links are misidentified mono-links

Among the intra-protein GVL-cross-links, we noticed that sometimes the two cross-linked peptides are adjacent to each other in the primary sequence. For example, the two peptides of the G-V cross-link in Figure 7A are aa265-aa274 (β-peptide) and aa275-aa293 (α-peptide) of the BSA protein. We wondered whether this BS^3^ cross-link might be a BS^3^ mono-link of aa265-aa293, because theoretically they have the same precursor mass and some of their fragment ions are also the same (the ones not containing a linked residue). Indeed, the MS2 of this G-V cross-link is better explained as a mono-link of aa265-aa293 with BS^3^ attached to K^274^ (Figure 7B). Likewise, the MS2 spectra of a subset of STY-cross-links are better explained by mono-links of the related, adjoined peptides. One such example is shown in Figure 7C-D.

**Figure 7.**
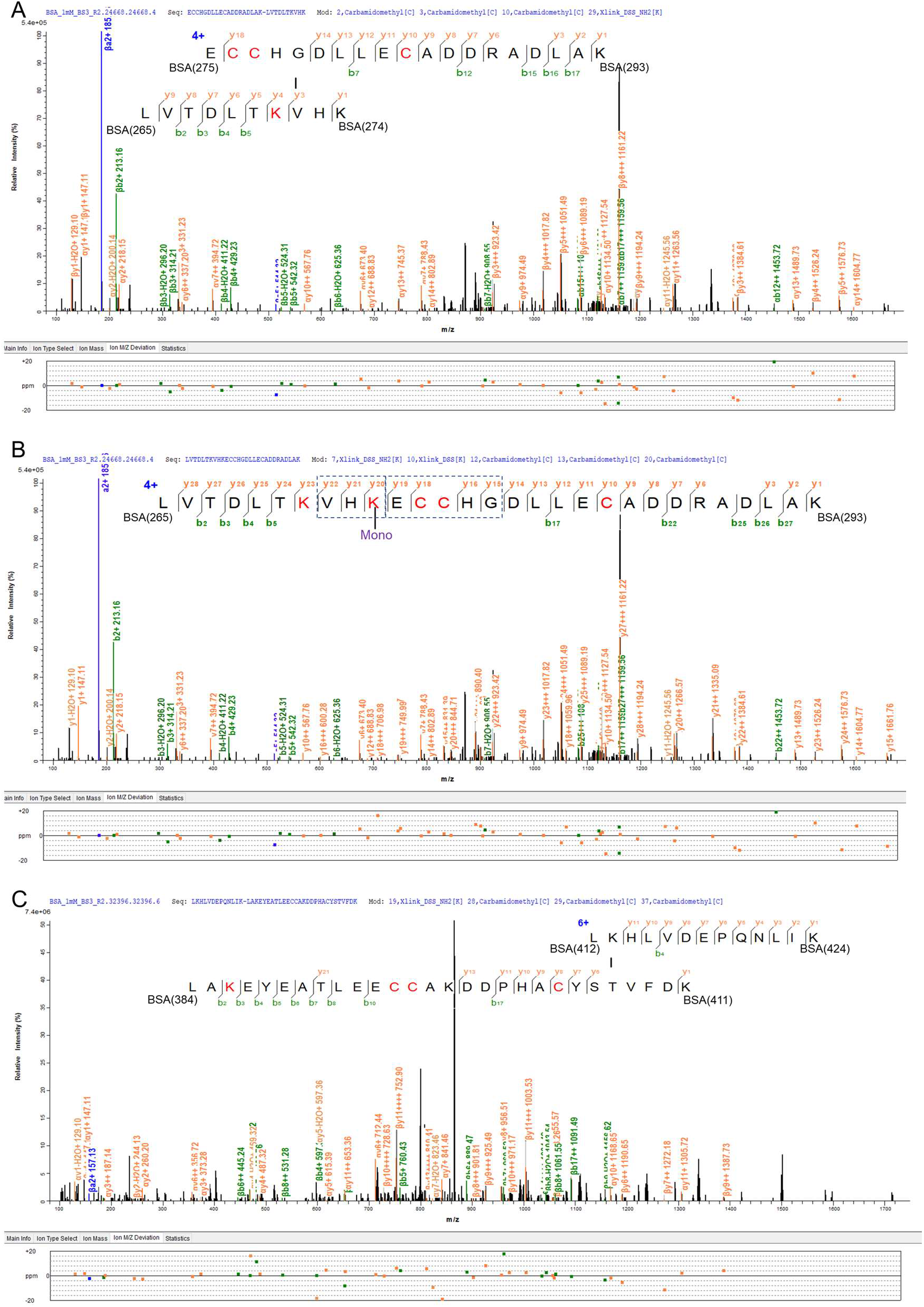
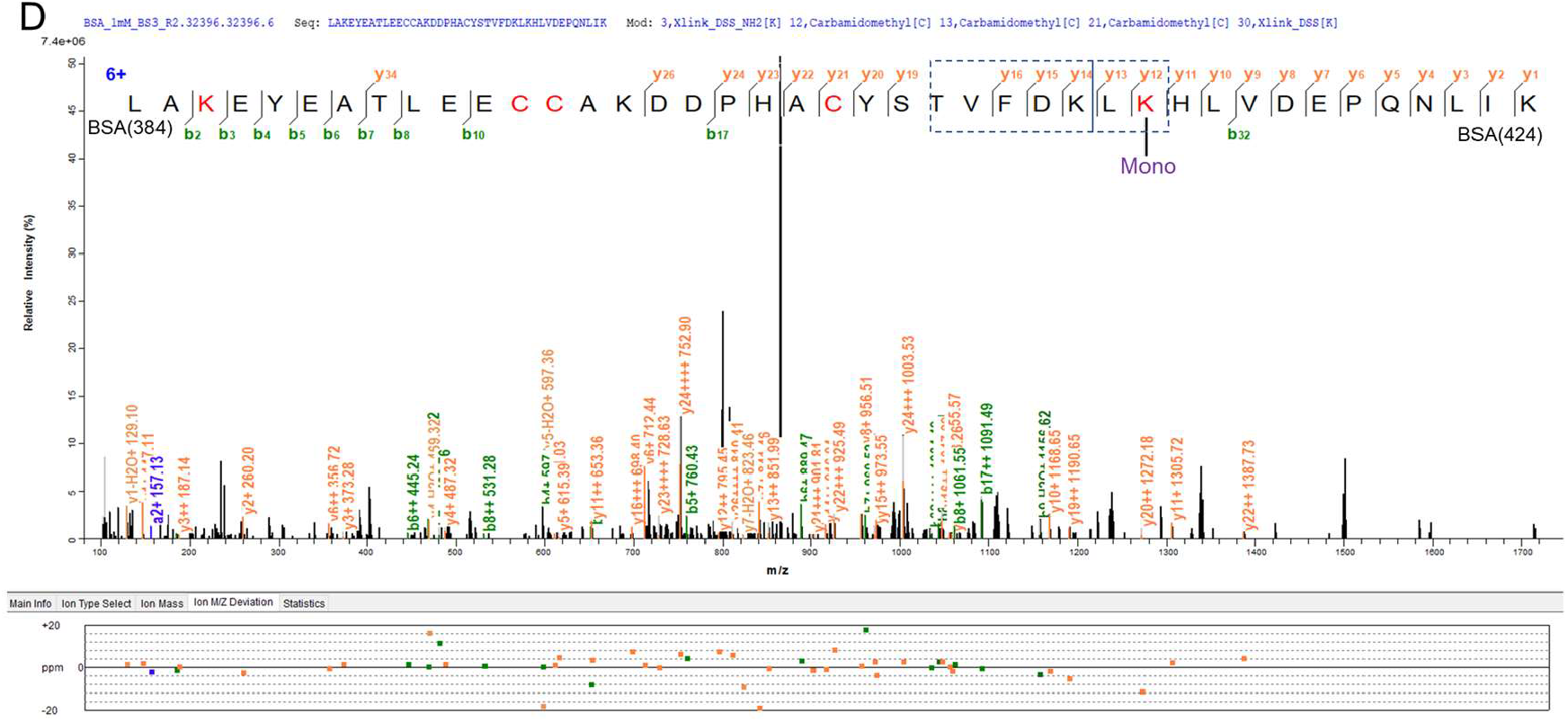
Two examples of K-mono-links misidentified as STY- or GVL-cross-links. (A-B) The same spectrum annotated as a G-V cross-link (A) or as a K-mono-link (B). (C-D) The same spectrum annotated as a K-T cross-link (C) or as a K-mono-link (D).

This type of false identification is not limited to BS^3^ or DSS, it is common to DSSO and DSBU as well (Figure 8). Although cross-linker independent, misidentification of mono-link as cross-link is search engine dependent; it occurs in xiSearch and pLink but not MeroX search results (Figure 8).The authors of MeroX have reported this type of misidentification^48^.

**Figure 8.**
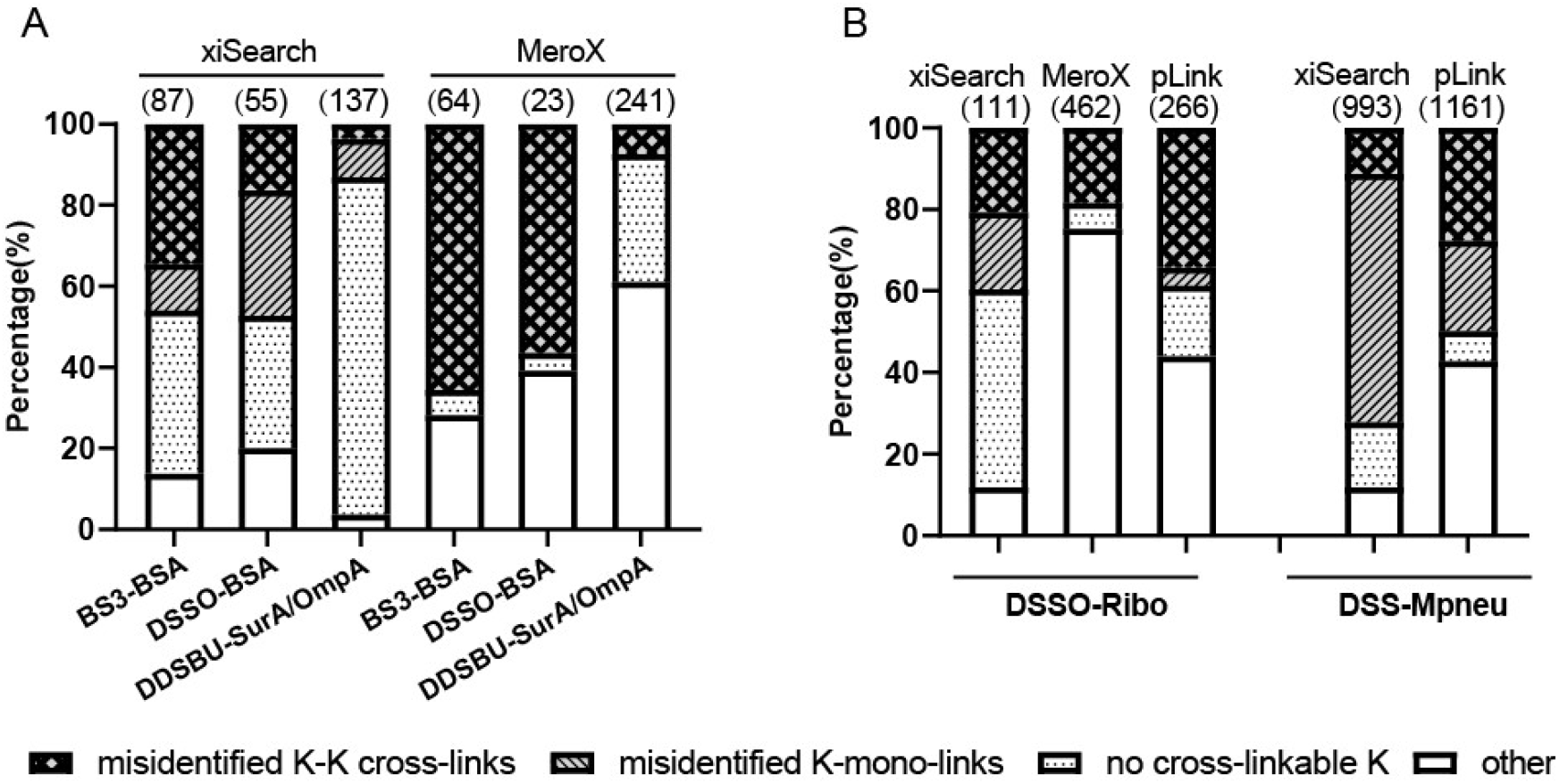
Summary of the origins of STY-cross-links. (A) low-complexity samples, three different cross-linkers. (B) medium- and high-complexity samples.

### An estimation of the contribution of different types of false identifications

In order to gauge the questionable cross-link identifications, we combined the STY-cross-links of the five datasets (DSS-BSA, DSSO-BSA, DSBU-SurA/OmpA, DSSO-Ribo, and DSS-Mpneu) and performed statistical analysis. xiSearch, MeroX and pLink identified a total of 1764, 578, and 1634 STY-cross-links, respectively. We find that 62.3%, 16.8% and 15.1% of them have no cross-linkable K in the peptide sequences (Supplementary Figure 5). For the rest, of the peptides having a link site on S, T, or Y, 50-75% have a cross-linkable K just 1-3 aa away from the reported link site. For these peptides, it is quite possible that the K-link sites are misidentified as STY-link sites. As the vast majority of STY-cross-links are K-STY cross-links, we estimate that close to 50-75% of the STY-crosslinks have a cross-linkable K 1-3 aa away from the reported link site, which means that the percentage of K-K cross-links misidentified as STY cross-links could be close to 18-64% of all STY-cross-links. This estimation is on par with the percentage of STY-cross-links that have a K-K cross-link “isoforms”, i.e., of the same two peptides but different link site(s) (Figure 8, segments marked by diamond pattern). For the MeroX search results on BS^3^-BSA and DSSO-BSA, this is a major source of misidentification, accounting for 57∼66% of the STY-cross-links. MeroX search results, however, have zero mono-links misidentified as STY-cross-links. For xiSearch and pLink, >5% of their STY-cross-links identifications are better explained as mono-links (Figure 8, slanted lines segments). On the DSS-Mpneu data, in particular, 61% of the STY-cross-links identified by xiSearch are misidentified mono-links. Similar results were obtained when the GVL-cross-links were analyzed (Supplementary Table 3-4).

For the other STY-cross-link identifications, a sizable portion contain no cross-linkable K. Analysis of this subset finds that 18-30% of them have at least on short peptide of 5-7 aa (Supplementary Figure 6). As seen from the benchmark dataset of synthetic peptides, the false, between-group cross-links have the same characteristic—about 50% of them have short peptides, compared with ∼30% for correct, within-group cross-links (Figure 4G-I). Identification of a cross-link involves identifying the sequence of two peptides and localizing link site in each. The above analysis shows that the three mainstream cross-link search engines are generally good at the former, but not so good at the latter task. Most, if not all, STY-cross-links are probably misidentifications and many of them are actually K-K cross-links. We thus recommend against setting STY as cross-linkable sites. A K-only cross-link search for NHS ester cross-linker increases accuracy without costing structural information gained from the data. As shown in Supplementary Table 1-2, if we ignore the difference in link sites, a K-only search and a KSTY search identify almost exactly the same number of peptide pairs form the cross-linked synthetic peptides (the number of identified mono-links in this standard dataset is close to zero).

## Discussion

Whether STY should be added as cross-linkable sites for NHS ester cross-linkers has been controversial for a long time. Based on the analysis of multiple CXMS datasets representing different NHS ester cross-linkers and different laboratories, and more importantly, with GVL-cross-link identifications as a negative control, we show that STY-cross-links should be rare in conditions commonly used for CXMS experiments, i.e., 0.2-2.0 mM NHS ester cross-linker, 25 ºC, 30-60 min for protein samples or up to h for peptide samples (Table I). Under such conditions, the STY- and GVL-cross-links identifications are not different in both the number and the quality score of CSMs. Therefore, most if not all STY-cross-link identification are unreliable. This is consistent with a recent finding that a pair of light/heavy isotope-labeled NHS ester probes did not label protein/ peptides at STY residue^46^. It is also supported by an earlier finding that at pH 6.7 and pH 7.8, no detectable reaction products were found between an NHS ester cross-linker and peptide hydroxyl groups of STY residues, even after 24 hours^30^. Therefore, setting STY as cross-linkable sites lacks justification from what is known about the chemical reactions of NHS esters in routine experimental settings.

**Table 1.**
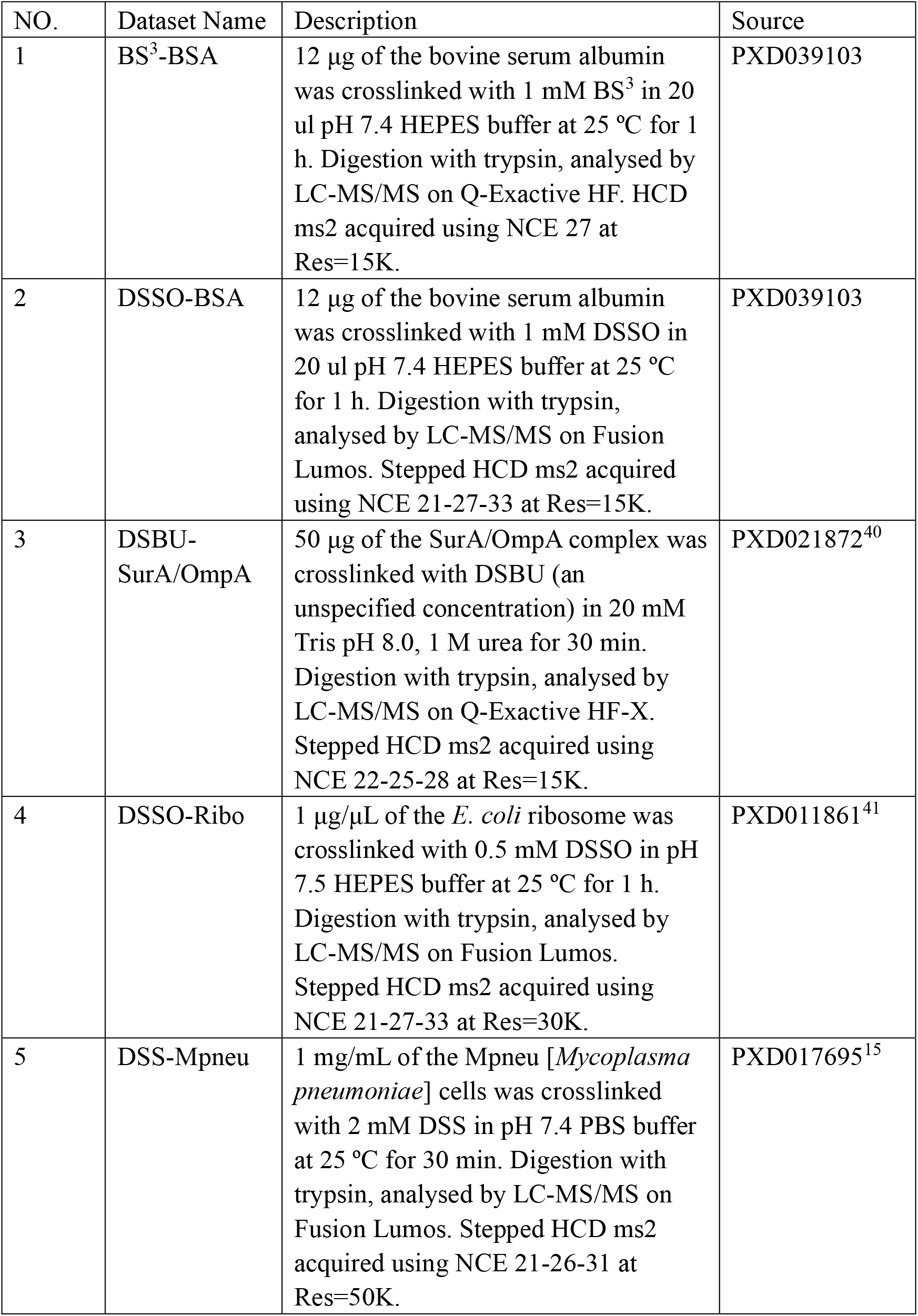
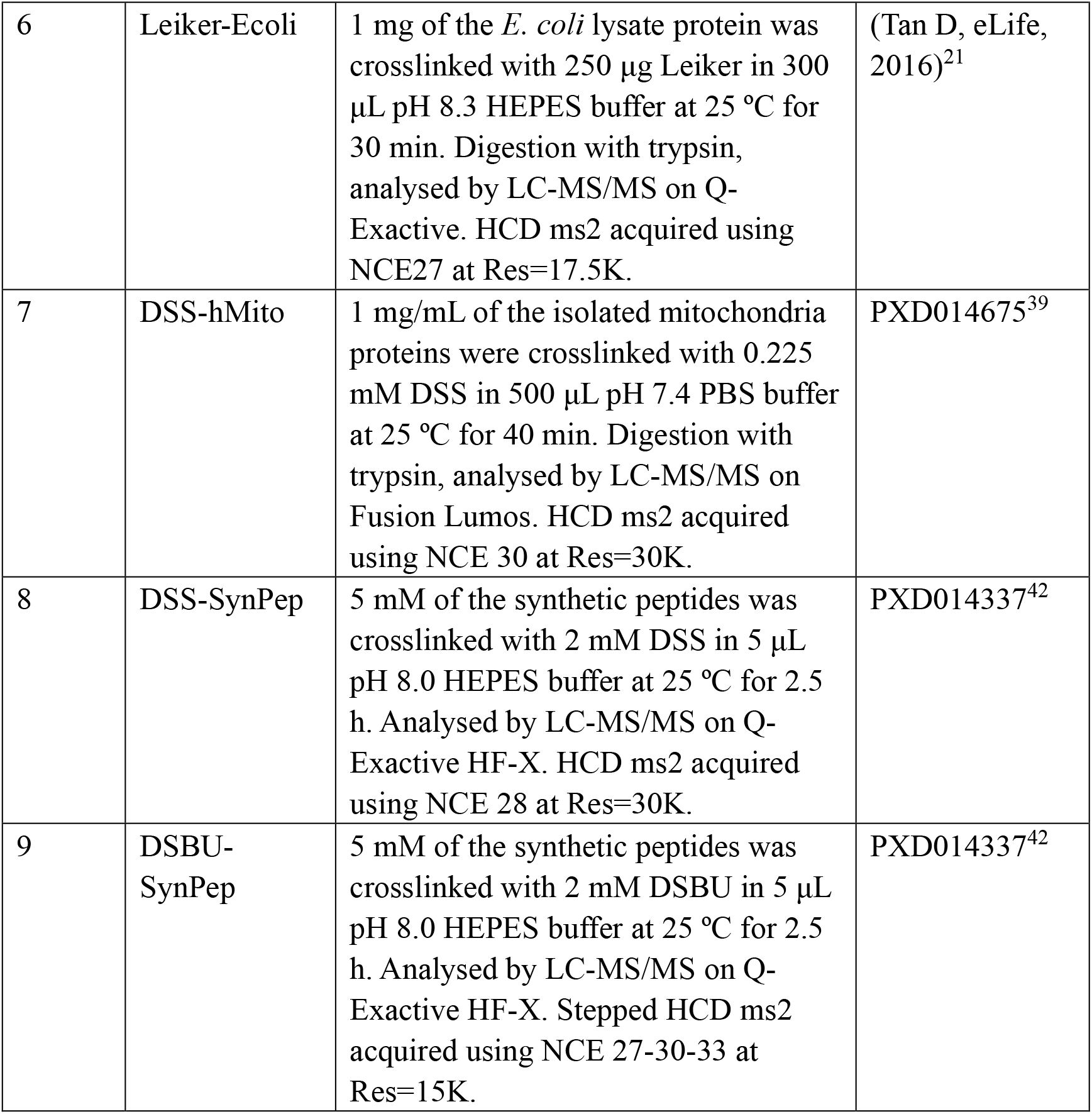
datasets used in this study

| NO. | Dataset Name | Description | Source |
| --- | --- | --- | --- |
| 1 | BS<sup>3</sup>-BSA | 12 µg of the bovine serum albumin was crosslinked with 1 mM BS<sup>3</sup> in 20 ul pH 7.4 HEPES buffer at 25 °C for 1 h. Digestion with trypsin, analysed by LC-MS/MS on Q-Exactive HF. HCD ms2 acquired using NCE 27 at Res=15K. | PXD039103 |
| 2 | DSSO-BSA | 12 µg of the bovine serum albumin was crosslinked with 1 mM DSSO in 20 ul pH 7.4 HEPES buffer at 25 °C for 1 h. Digestion with trypsin, analysed by LC-MS/MS on Fusion Lumos. Stepped HCD ms2 acquired using NCE 21-27-33 at Res=15K. | PXD039103 |
| 3 | DSBU-SurA/OmpA | 50 µg of the SurA/OmpA complex was crosslinked with DSBU (an unspecified concentration) in 20 mM Tris pH 8.0, 1 M urea for 30 min. Digestion with trypsin, analysed by LC-MS/MS on Q-Exactive HF-X. Stepped HCD ms2 acquired using NCE 22-25-28 at Res=15K. | PXD021872<sup>40</sup> |
| 4 | DSSO-Ribo | 1 µg/µL of the *E. coli* ribosome was crosslinked with 0.5 mM DSSO in pH 7.5 HEPES buffer at 25 °C for 1 h. Digestion with trypsin, analysed by LC-MS/MS on Fusion Lumos. Stepped HCD ms2 acquired using NCE 21-27-33 at Res=30K. | PXD011861<sup>41</sup> |
| 5 | DSS-Mpneu | 1 mg/mL of the Mpneu [*Mycoplasma pneumoniae*] cells was crosslinked with 2 mM DSS in pH 7.4 PBS buffer at 25 °C for 30 min. Digestion with trypsin, analysed by LC-MS/MS on Fusion Lumos. Stepped HCD ms2 acquired using NCE 21-26-31 at Res=50K. | PXD017695<sup>15</sup> |
| 6 | Leiker-Ecoli | 1 mg of the *E. coli* lysate protein was crosslinked with 250 µg Leiker in 300 µL pH 8.3 HEPES buffer at 25 °C for 30 min. Digestion with trypsin, analysed by LC-MS/MS on Q-Exactive. HCD ms2 acquired using NCE27 at Res=17.5K. | (Tan D, eLife, 2016)<sup>21</sup> |
| 7 | DSS-hMito | 1 mg/mL of the isolated mitochondria proteins were crosslinked with 0.225 mM DSS in 500 µL pH 7.4 PBS buffer at 25 °C for 40 min. Digestion with trypsin, analysed by LC-MS/MS on Fusion Lumos. HCD ms2 acquired using NCE 30 at Res=30K. | PXD014675<sup>39</sup> |
| 8 | DSS-SynPep | 5 mM of the synthetic peptides was crosslinked with 2 mM DSS in 5 µL pH 8.0 HEPES buffer at 25 °C for 2.5 h. Analysed by LC-MS/MS on Q-Exactive HF-X. HCD ms2 acquired using NCE 28 at Res=30K. | PXD014337<sup>42</sup> |
| 9 | DSBU-SynPep | 5 mM of the synthetic peptides was crosslinked with 2 mM DSBU in 5 µL pH 8.0 HEPES buffer at 25 °C for 2.5 h. Analysed by LC-MS/MS on Q-Exactive HF-X. Stepped HCD ms2 acquired using NCE 27-30-33 at Res=15K. | PXD014337<sup>42</sup> |

So far, the evidence that supports STY-cross-links comes from incidents of good-looking spectral match. For example, Ryl et al. showed a well-matched spectrum to prove that serine could be cross-linked^39^. As we show in this study, although many GVL-cross-link identifications have poorly-matched spectra, some of them do have surprisingly good-looking spectra (Supplementary Figure 7B-C). This goes to show that an occasionally seen good match or two are insufficient evidence to prove that certain cross-links do occur. When the sequences of the two peptides are identified correctly, the appearance or disappearance of a tiny peak can shift the link site assignment from one residue to another (Supplementary Figure 4C-D). Without additional information, such as the knowledge of cross-linker specificity, precise localization of the link site needs contiguous fragment ions and the presence of isotopic peaks to validate their identities (charge state and monoisotopic *m/z*).

The three categories of STY-cross-links—(I) misidentified K-K cross-links, (II) misidentified mono-links, and (III) no cross-linkable K—have different consequences on structural interpretation of the CXMS data. For K-K cross-links misidentified as STY-cross-links, the sequences of the two linked peptides are correctly identified, the link sites not. As such, the structural interpretation that a category I STY-cross-link will lead to is not entirely wrong, but imprecise. For mono-links misidentified as STY-cross-links, they are not so informative to begin with because the two peptides are adjacent to each other in primary sequence of a protein. In other words, they have limited negative consequences. For the last category of STY-cross-links, 4.3-87.2% have no cross-linkable lysine residues and they tend to have a short peptide of ≤ 7 aa 1.5-30.7%, which is a feature shared by the confirmed false identifications in the grouped synthetic peptide dataset. This suggests that many of the category III STY-cross-links likely have incorrect peptide sequence identifications, let alone link-site assignment, they will surely mislead structural interpretation and cause more damage than the Category I and Category II cross-links.

Adding STY or GVL as cross-linkable sites increases the search space for peptide/cross-link-spectrum matching. Even if FDR is controlled at an acceptable level, the expanded base can lead to an increase in the number of falsely identified PSMs or CSMs. An examination of the cross-link search space, namely, the number of possible combinations of peptide pairs in a link-site-sensitive manner seems to agree with this idea. As shown in Figures 1-2 and Supplementary Figure 1, the search space (red squares. Y-axis on the right) of the nonK-nonK cross-links is often one-order of magnitude greater than that of K-K cross-links. A similar pattern is seen for mono-link search (Figure 3). Considering the large search space, the current cross-link search engines have done a good job of keeping out the vast majority of true negatives, namely, all GVL-cross-links and by the deduction above, most if not all STY-cross-links.

In summary, a weakness shared by the cross-link search engines has been found in this study. For a small but significant fraction of the CSMs that pass the filtering criteria, the link sites are not located precisely. This can become a serious problem when multiple amino acids are set as cross-linkable sites, and it is a great challenge to be faced by cross-link search engines when analyzing CXMS data of photoactivated cross-linkers, since they could theoretically react with any amino acid residues.

## Acknowledgements

We thank Prof. Si-Min He and Prof. Hao Chi for their advice in experiment design and manuscript revision. The authors gratefully acknowledge financial support from the Ministry of Science and Technology of China (2020YFF01014505 to M.-Q.D.), the municipal government of Beijing (in the form of NIBS intramural grants), TIMBR, and Tsinghua University.

## Author Contributions

M.-Q.D. devised the project. Y.C. and M.-Q.D. designed the experiments in this study. Y.C. performed the most data analysis. X.-T.L. performed partial data analysis of pLink. P.-Z.M. performed the ^15^N-MS1 evaluation of identified cross-links of xiSearch, MeroX, and pLink. C.T. developed a unility to visualize the elution peak of cross-linked precursors. Y.C. and M.-Q.D. wrote the manuscript.

## Note

The authors declare no competing financial interest.

