## Supplementary Information for "Comparative analysis of chemical cross-linking mass spectrometry data indicates that protein STY residues rarely react with N-hydroxysuccinimide ester cross-linkers"

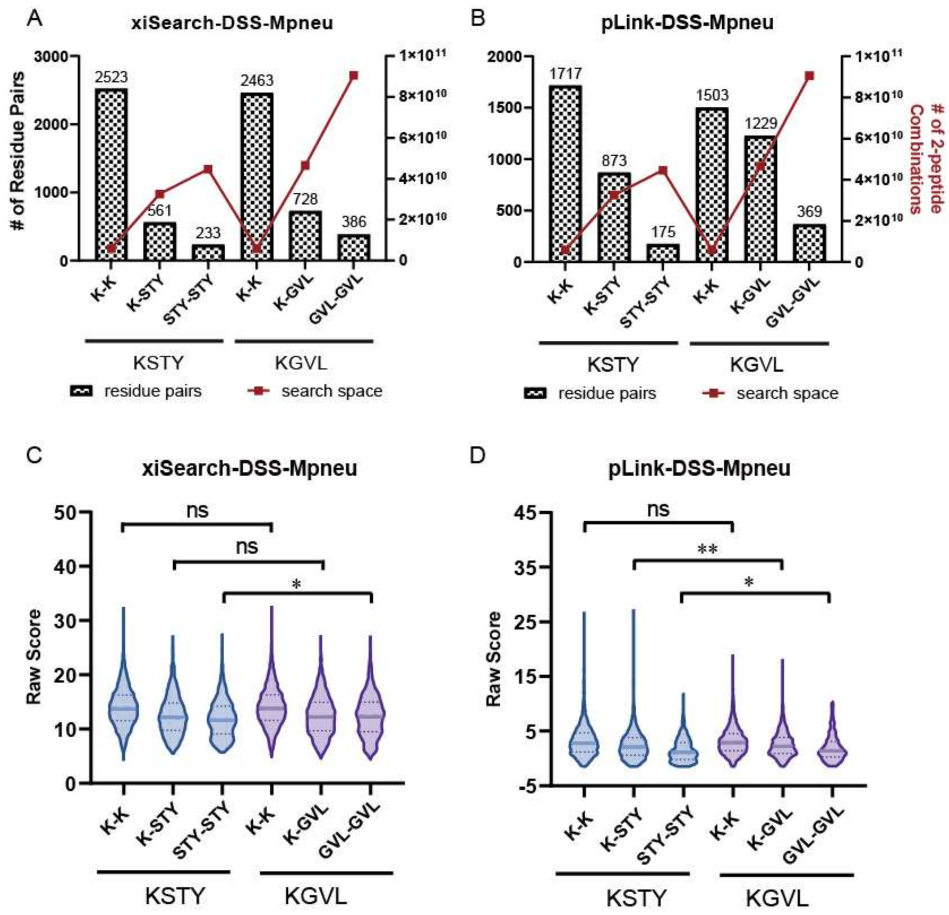

**Supplementary Figure 1.** STY-cross-links are only as reliable as GVL-cross-links on high-complexity sample regardless of the search engines used to identify them.

The numbers (A-B) and the CSM score distributions (C-D) of the STY- and GVL-cross-link identifications by xiSearch and pLink on the high-complexity DSS-Mpneu dataset. In (A-C) the number of identified residue pair are indicated by bars (left y-axis) and the size of the search space as measured by the numbers of possible peptide pair combinations is indicated by line-connected red squares (right y-axis).

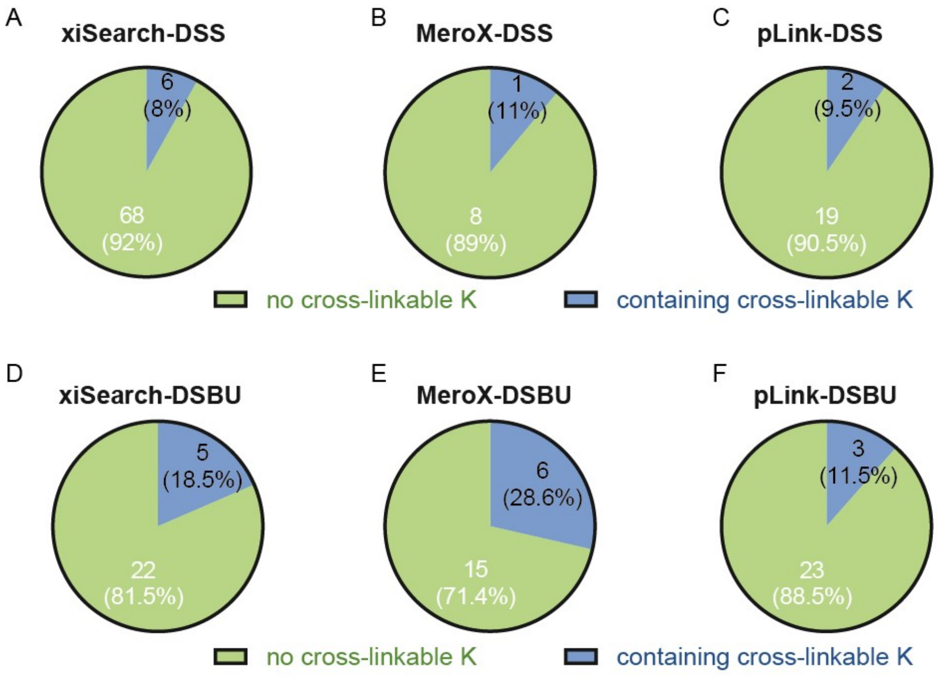

**Supplementary Figure 2.** analysis of the mistaken STY-cross-linked peptide pairs in Figure 4.
(A-C) the percentage of peptide pairs that contain at least one peptide without cross-linkable lysine in the mistaken peptides on the DSS-SynPep dataset by (A-C) or the DSBU-SynPep dataset (D-F) by xiSearch, MeroX, and pLink.
Note: a cross-linkable K does not include the C-terminal lysine produced by tryptic digestion. Once modified by a NHS ester, K becomes resistant to trypsin digestion.

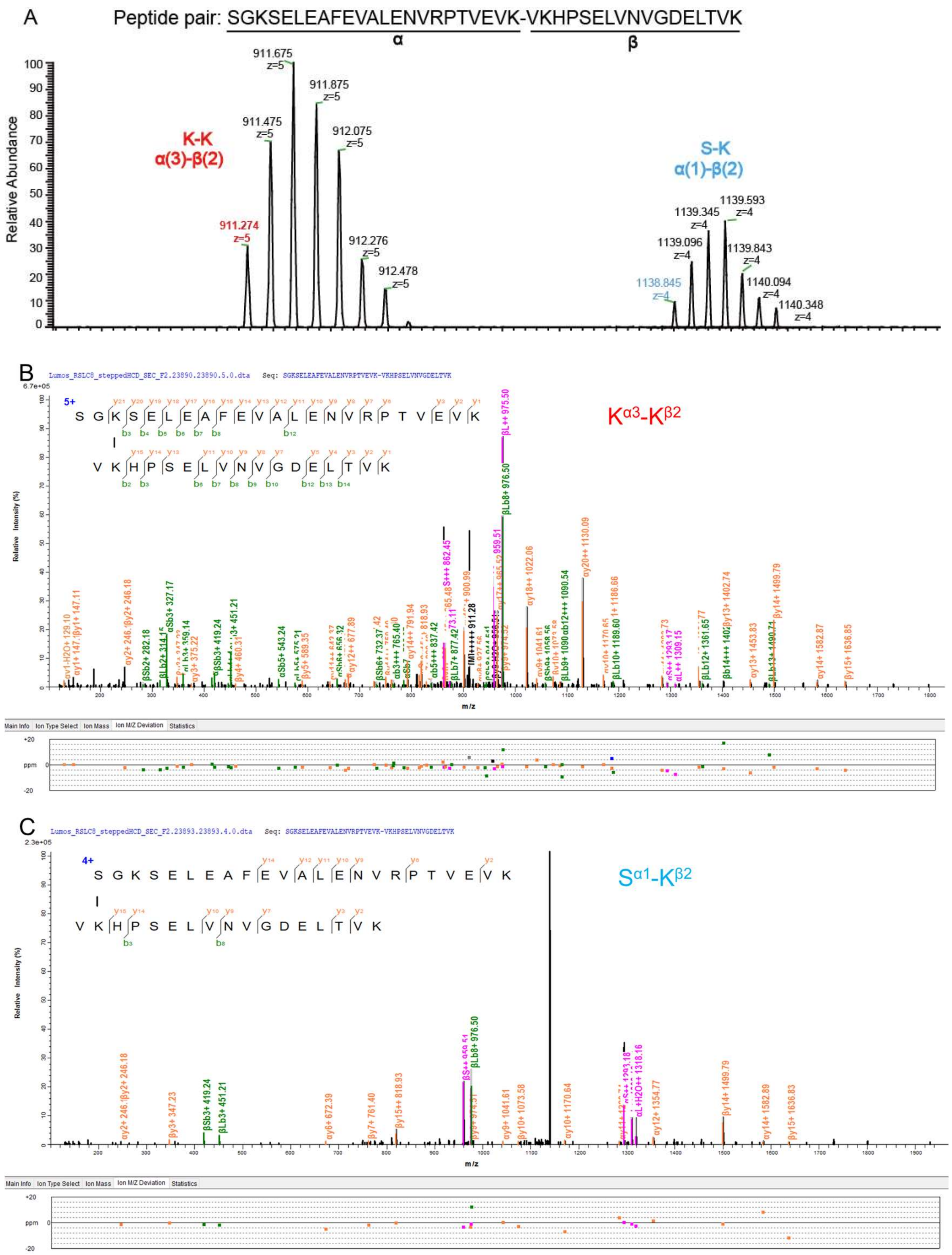

**Supplementary Figure 3.** K-K cross-links misidentified as STY-cross-links.

(A) The isotopic peak of $4^{+}$, $5^{+}$ precursor of cross-linked peptide pair (SGKSELEAFEVALENVRPTVEVK-VKHPSELVNVGDELTVK), the following MS2 spectra were identified with different linkage. (B) The cross-link-spectrum-match of K-K linkage α(3)-β(2). (C) The cross-link -spectrum-match of the S-K linkage α(1)-β(2).

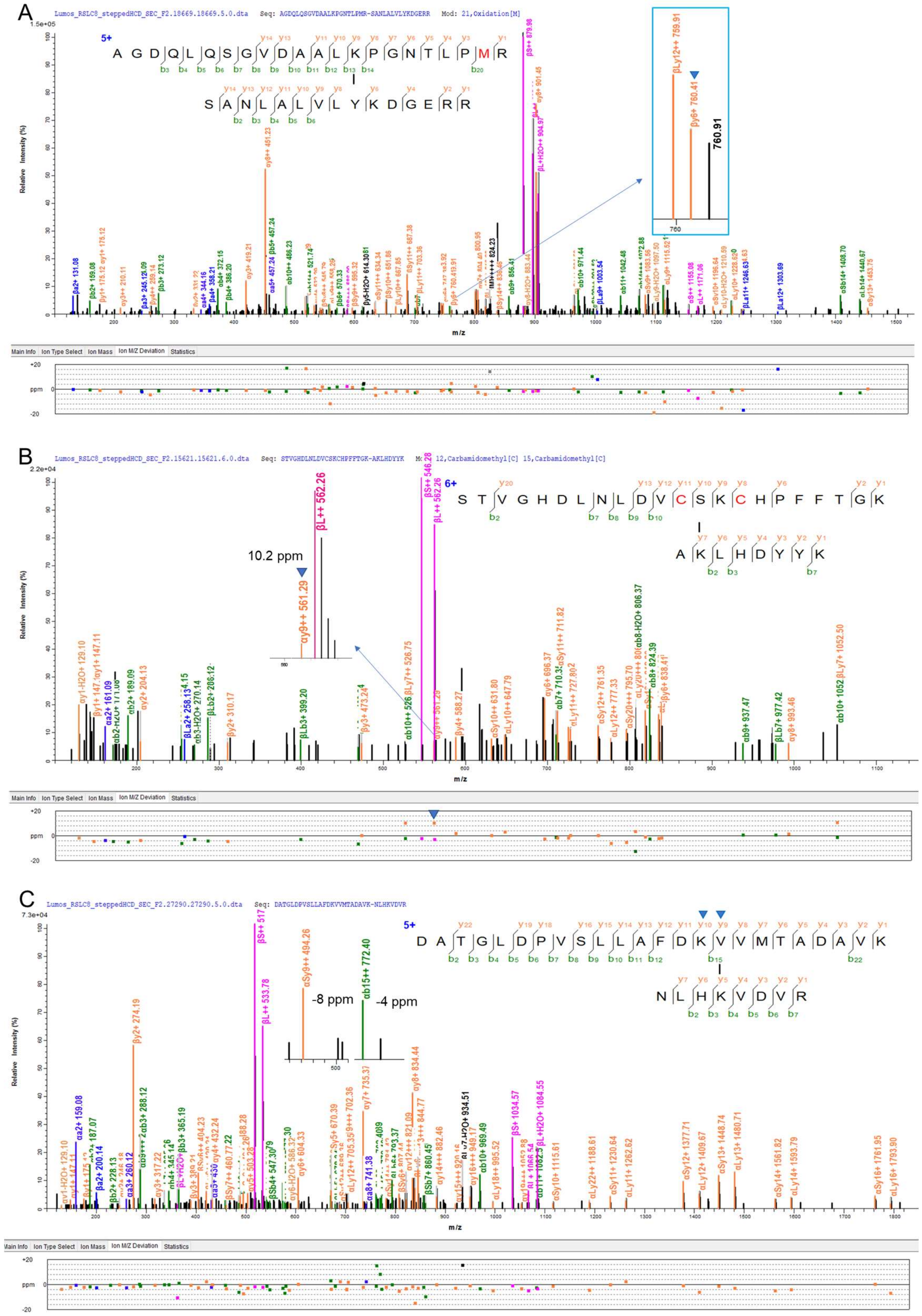

A
Lumos_RSLC8_steppedHCD_SEC_F2.18669.18669.5.0.dta Seq: AGDQLQSGVDAALKPGNTLPMR-SANLALVLYKDGERR Mod: 21,Oxidation[M]
Relative Intensity (%)
m/z
760.91
B
Lumos_RSLC8_steppedHCD_SEC_F2.15621.15621.6.0.dta Seq: STVGHDLNLDVCSKCHPFFTGK-AKLHDYYK Mod: 12,Carbamidomethyl[C] 15,Carbamidomethyl[C]
10.2 ppm
Relative Intensity (%)
m/z
C
Lumos_RSLC8_steppedHCD_SEC_F2.27290.27290.5.0.dta Seq: DATGLDPVSLLAFDKVVMTADAVK-NLHKVDVR
-8 ppm
-4 ppm
Relative Intensity (%)
m/z

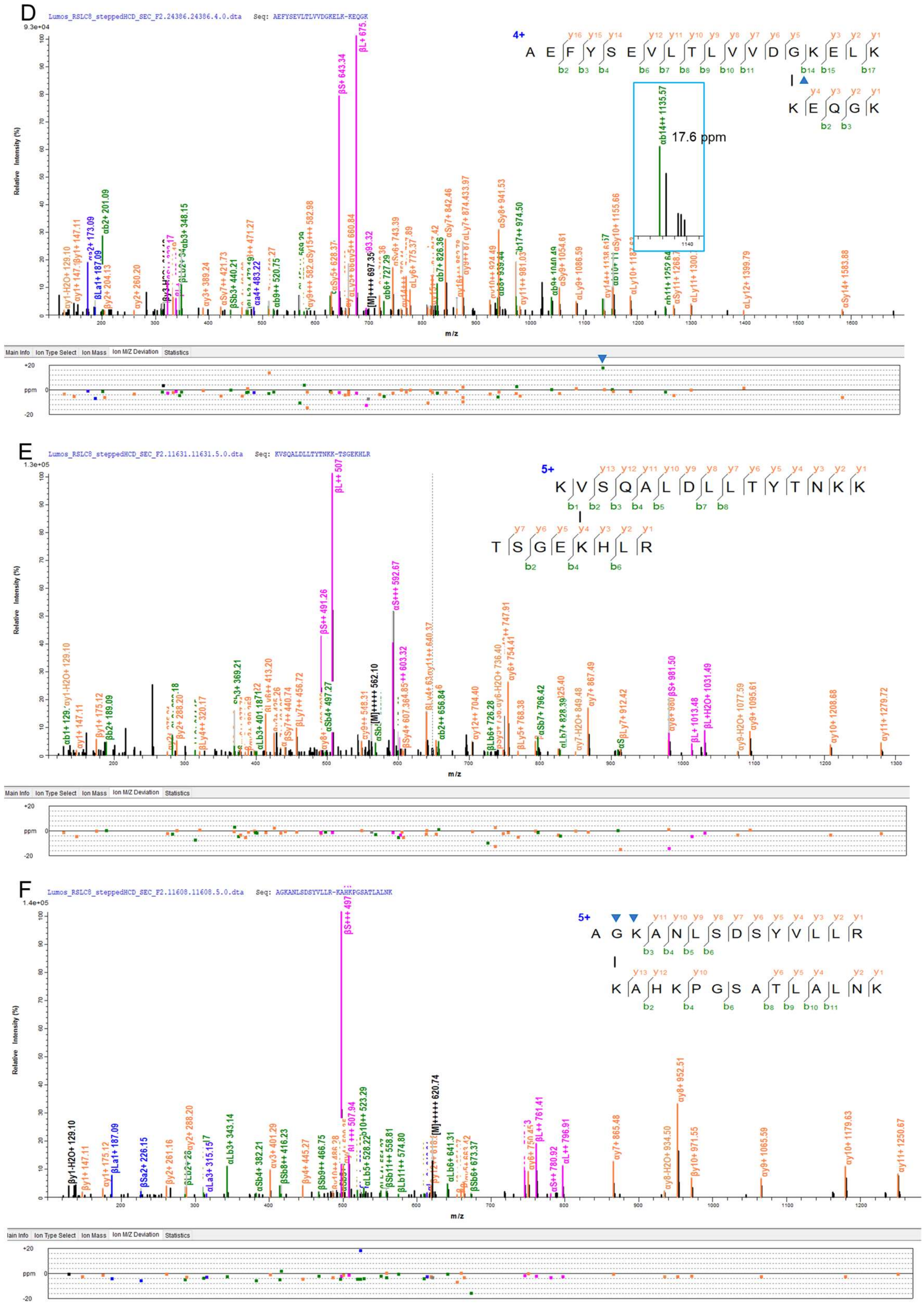
D
Lumos_RSLC8_steppedHCD_SEC_F2.24386.24386.4.0.dta Seq: AEFYSEVLTLVVDGKELK-KEQGK
4+
A E F Y S E V L T L V V D G K E L K
K E Q G K
17.6 ppm
Relative Intensity (%)
m/z
Main Info
Ion Type Select
Ion Mass
Ion M/Z Deviation
Statistics
ppm
E
Lumos_RSLC8_steppedHCD_SEC_F2.11631.11631.5.0.dta Seq: KVSQALDLLTYTNKK-TSGEKHLR
5+
K V S Q A L D L L T Y T N K K
T S G E K H L R
Relative Intensity (%)
m/z
F
Lumos_RSLC8_steppedHCD_SEC_F2.11608.11608.5.0.dta Seq: AGKANLSDSYVLLR-KAHKPGSATLALNK
5+
A G K A N L S D S Y V L L R
K A H K P G S A T L A L N K
Relative Intensity (%)
m/z

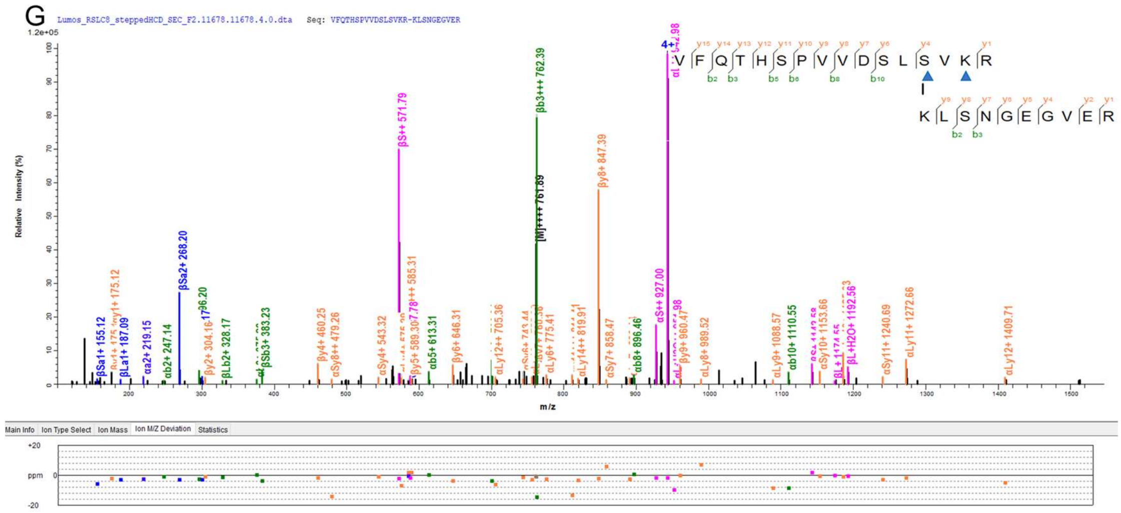

**Supplementary Figure 4.** Examples that the STY- or GVL-cross-links might be mislocated.

(A) a spectrum of K-Y cross-linking. The only evidence for positioning the link site at $Y^{\beta 9}$, not $K^{\beta 10}$ is an ion at m/z 760.41, which is interpreted by the cross-link search engine as $\beta y_6^+$. If this is true, then $\beta y_6^+$ has a lone monoisotopic peak in the MS2 spectrum. Unaccompanied by any isotopic peaks (761.41, 762.41). A better interpretation is the first of the two isotopic peaks of $\beta Ly_{12}^{++}$, in a three-peak cluster consisting of m/z 759.91, 760.41, and 760.91 (inset of Supplementary Figure 3A). In summary, upon close inspection, there is actually no evidence for $Y^{\beta 9}$ being the link site, $K^{\beta 10}$, the lysine residue adjacent to $Y^{\beta 9}$ should be the link site.

(B) a spectrum with K-S cross-linking. An apparent $\alpha y_9^{++}$ at m/z 561.29 is the only evidence for positioning the link site at $S^{\alpha 13}$ instead of $K^{\alpha 14}$. This $\alpha y_9^{++}$ is a low-intensity peak with no isotopic peaks to verity its 2+ charge state. Alarmingly, it has a mass deviation of 10 ppm, while most other fragment ions have a mass deviation of -7 ppm to 3 ppm. Therefore, localization of the link site to $S^{\alpha 13}$ instead of $K^{\alpha 14}$ is again based on the questionable evidence of one tiny peak.

(C) a well-matched spectrum of K-G cross-linking, whose mass peak supported α16(V) is link site, but not adjacent α15(K). This spectrum demonstrates that chance match can sometimes be deceptive enough to derail the link site identification. There are many peaks in this MS2 spectrum, especially below m/z 900. For the link site on α-peptide, $V^{\alpha 16}$ is chosen over $K^{\alpha 15}$ due to the presence of two ions: m/z 494.26 happens to be a match for $\alpha Sy_9^{++}$ and thus supports $V^{\alpha 16}$ as link site; m/z 772.40 happens to be a match for $\alpha b_{15}^{++}$ and thus argue against $K^{\alpha 15}$ being the link site.

(D) a spectrum of K-G cross-linking. The only evidence for $G^{\alpha 14}$ being the link site instead of the adjacent $K^{\alpha 15}$ is m/z 1135.57, which matched the theoretical mass of $\alpha b_{14}^{++}$ if it's a doubly charged ion. This is an unreliable interpretation for the following reasons. First, there are no isotopic peaks to verify that m/z 1135.57 is a doubly charged ions. Second, if it were doubly charged, it's mass of ~2270 Da would predict that the first isotopic peak (m/z 1136.07) should have a higher intensity and thus more readily detectable than the monoisotopic peak (m/z 1135.57). However, m/z 1136.07 is absent in the spectrum. Third, this purported $\alpha b_{14}^{++}$ fragment has a mass deviation of >16 ppm,

markedly above other fragment ions (mostly -8 ppm to 4 ppm).
(E) a spectrum of impossible K-V cross-linking. An apparent $\alpha b_1^{1+}$ at m/z 129.10 is the only fragment ion that supports $V^{\alpha2}$ (the superscript denotes the peptide and the amino acid position, α2 is the second amino acid of the α peptide) not $K^{\alpha1}$, being the link site. However, the fragmentation study in the past have shown that $b_2$ is the smallest member in the b series. For this reason and that the α peptide happens to have a K at C-terminus, the most logical explanation for the m/z 129.10 $\alpha y^{1+}$-$H_2O$, not $\alpha b_1^{1+}$. In other words, there's no convincing evidence for $V^{\alpha2}$ being the link site, the link site should be $K^{\alpha1}$.
(F) a spectrum of K-G cross-linking. No cleavage is detected between Gα2, the software assigned link site, and Kα3. In fact, the fragment ions in this MS2 spectrum can only marrow down the link site on the α-peptide to the first three residues (AGK), and only K is chemically valid link site.
(G) a spectrum with K-S cross-linking. Assigning the link site to Sα13 instead of the adjacent Kα15 is clearly by chance, not by evidence, because fragment ions resulting from a cleavage between Sα13 and Kα15 are not seen in the spectrum.

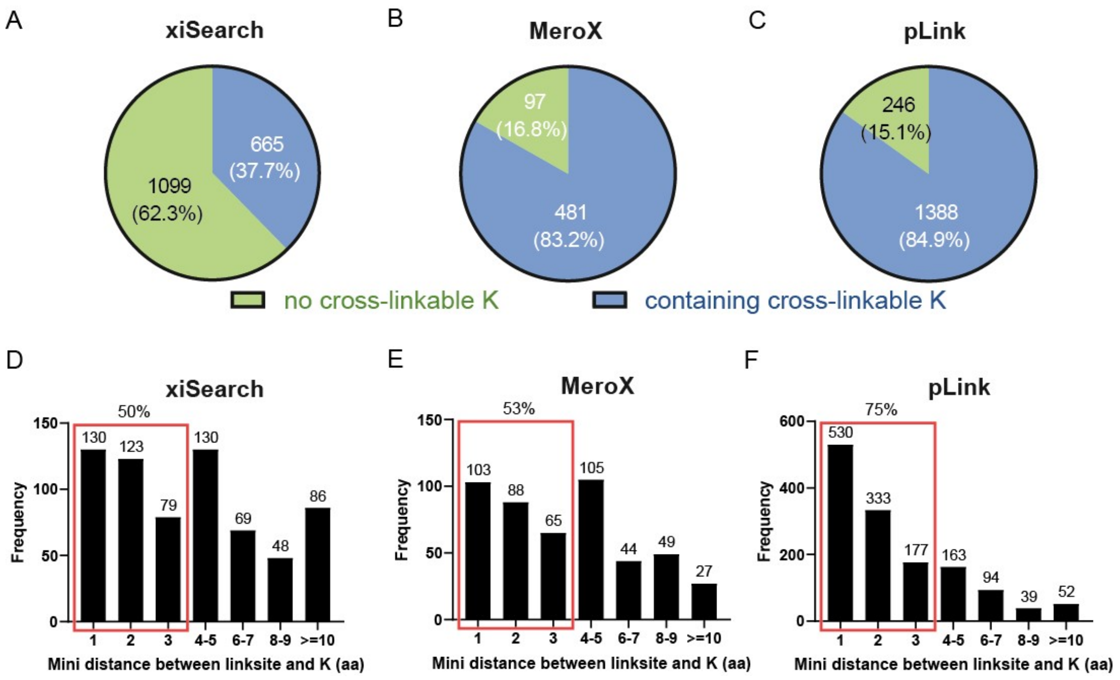

**Supplementary Figure 5.** Link-site localization may be imprecise for many STY-cross-links.

(A-C) Percentage of STY-cross-links with or without a cross-linkable lysine in cis with the STY link site.

Note: a cross-linkable K does not include the C-terminal lysine produced by tryptic digestion. Once modified by a NHS ester, K becomes resistant to trypsin digestion.

(D-F) The distribution of the minimal distance (aa) between the identified STY link site and a cross-linkable lysine by (D) xiSearch, (E) MeroX, and (F) pLink search results.

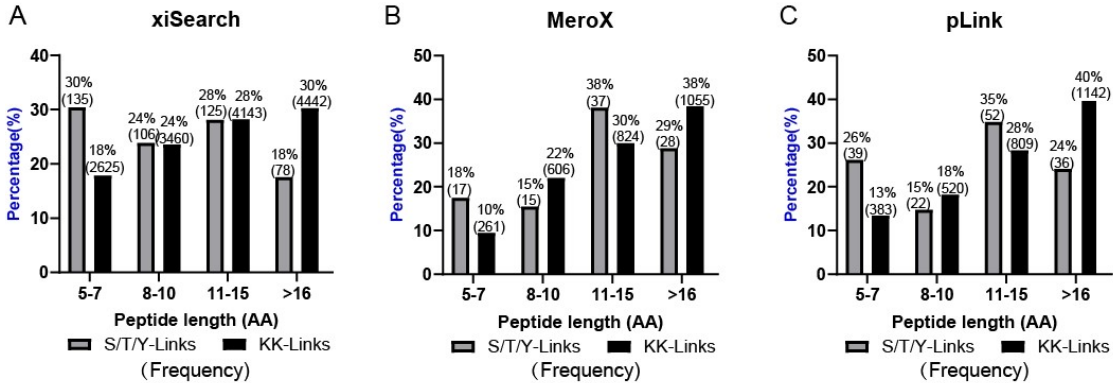

**Supplementary Figure 6.** STY-cross-links that are not readily accounted for by K-mono-links or K-K cross-links and are devoid of cross-linkable K tend to contain short peptides of 5-7 aa. K-K cross-links identified by the same search engine serve as a control. The datasets in Figure 1-2 and Supplementary Figure 1 are used in this analysis. (A) xiSearch, (B) MeroX, (C) pLink.

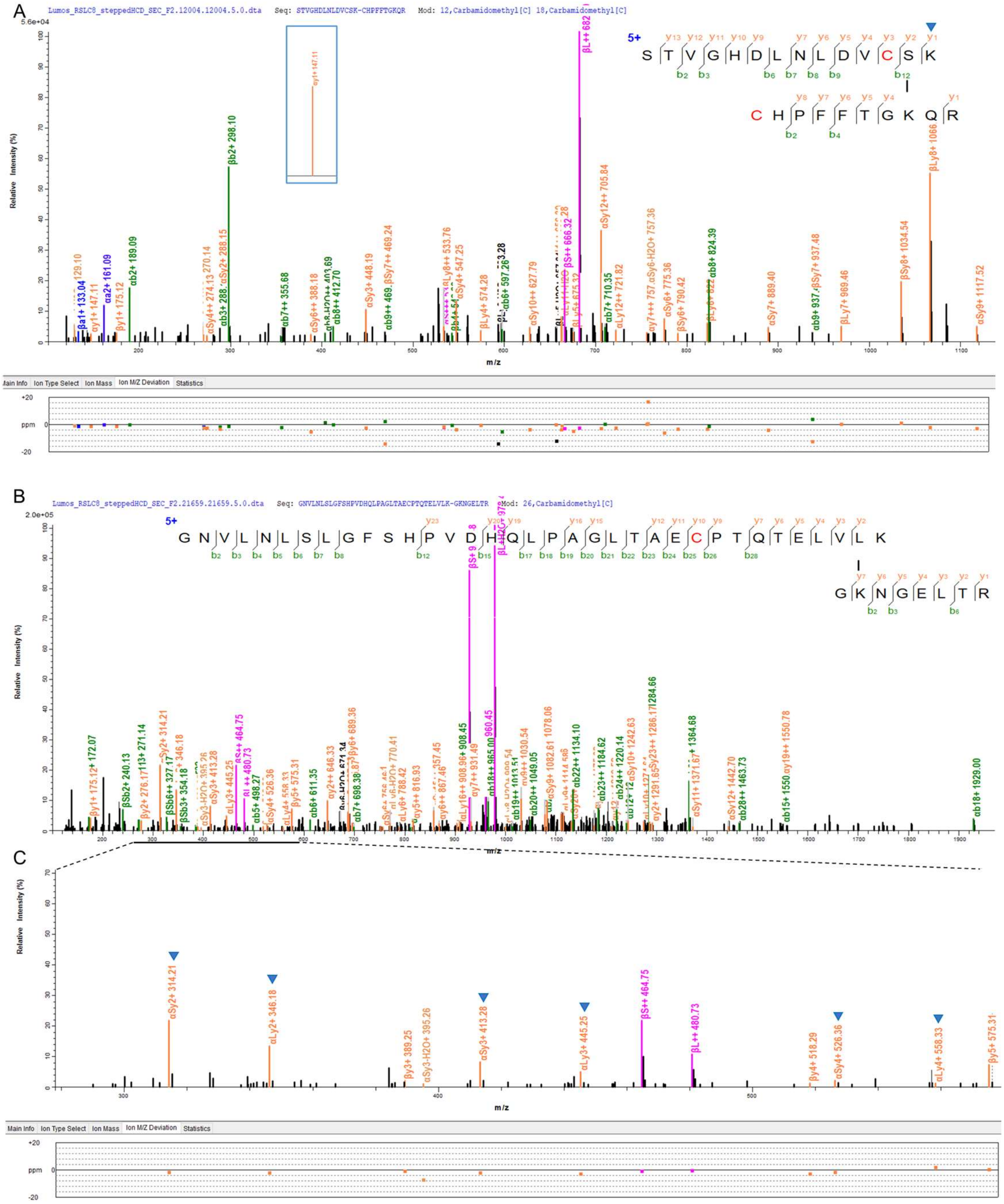

**Supplementary Figure 7.** Two high-quality spectra of STY or GVL could be cross-links.

(A) a well-matched spectrum of a K-S cross-link. (B) a well-matched spectrum of a K-L cross-link. (C) the zoom view of m/z 250~600 in C, the mass peaks $\alpha Sy_2^+$, $\alpha Ly_2^+$, $\alpha Sy_3^+$, $\alpha Ly_3^+$, $\alpha Sy_4^+$, $\alpha Ly_4^+$ that support link site is L, are labeled by the blue triangle.

**Supplementary Table 1.** the number of peptide pair and sequence pair of the correct identification in Figure 4A-C.

| | xiSearch | | MeroX | | pLink | |
|---|---|---|---|---|---|---|
| | KSTY-KSTY | K-K | K-KSTY | K-K | KSTY-KSTY | K-K |
| Peptide pair with site | 327 | 333 | 106 | 87 | 334 | 243 |
| sequence pair | 247 | 254 | 88 | 87 | 238 | 243 |

**Supplementary Table 2.** the number of peptide pair and sequence pair of the correct identification in Figure 4D-F.

| | xiSearch | | MeroX | | pLink | |
|---|---|---|---|---|---|---|
| | KSTY-KSTY | K-K | K-KSTY | K-K | KSTY-KSTY | K-K |
| Peptide pair with site | 343 | 339 | 413 | 326 | 318 | 268 |
| sequence pair | 250 | 270 | 324 | 326 | 267 | 268 |

**Supplementary Table 3.** The percentage of STY(GVL)-linked peptide pairs that have the corresponding KK pairs on five datasets.

| | xiSearch | |
|---|---|---|
| Dataset | STY-links | GVL-links |
| $BS^3$-BSA | 34.5% (30/87) | 19.6% (22/112) |
| DSSO-BSA | 16.4% (9/55) | 20.6% (22/107) |
| DSBU-SurA/OmpA | 3.6% (5/137) | 1.4% (2/144) |
| DSSO-Ribo | 20.7% (23/111) | 27.7% (56/202) |
| DSS-Mpneu | 11.3% (112/993) | 11.8% (165/1402) |
| | pLink | |
| | STY-links | GVL-links |
| DSSO-Ribo | 34.2% (91/266) | 32.8% (213/649) |
| DSS-Mpneu | 27.6% (321/1161) | 27.1% (481/1773) |
| | MeroX | |
| | STY-links | GVL-links |
| $BS^3$-BSA | 65.6% (42/64) | 65.1% (69/106) |
| DSSO-BSA | 56.5% (13/23) | 45.0% (9/20) |
| DSBU-SurA/OmpA | 7.5% (18/241) | 6.2% (16/258) |
| DSSO-Ribo | 18.4% (85/462) | 18.8% (47/250) |

Supplementary Table 4. The percentage of STY(GVL)-linked peptide pairs that were most likely mono-links (end-to-end in protein sequence) on five datasets.

| | xiSearch | |
|---|---|---|
| Dataset | STY-links | GVL-links |
| $BS^3$-BSA | 11.5% (10/87) | 29.5% (33/112) |
| DSSO-BSA | 30.9% (17/55) | 42.1% (45/107) |
| DSBU-SurA/OmpA | 9.5% (13/137) | 13.9% (20/144) |
| DSSO-Ribo | 18.9% (21/111) | 27.2% (55/202) |
| DSS-Mpneu | 60.9% (605/993) | 64.1% (899/1402) |
| | pLink | |
| | STY-links | GVL-links |
| DSSO-Ribo | 4.5% (12/266) | 6.9% (45/649) |
| DSS-Mpneu | 22.2% (258/1161) | 24.9% (442/1773) |
| | MeroX | |
| | STY-links | GVL-links |
| $BS^3$-BSA | 0.0% (0/64) | 0.0% (0/106) |
| DSSO-BSA | 0.0% (0/23) | 0.0% (0/20) |
| DSBU-SurA/OmpA | 0.0% (0/241) | 0.0% (0/258) |
| DSSO-Ribo | 0.0% (0/462) | 0.0% (0/250) |
